# The inositol pyrophosphate 5-InsP7 promotes DNA repair by disrupting RAD51 binding to the C-terminus of BRCA2

**DOI:** 10.64898/2026.05.10.723974

**Authors:** Shubhra Ganguli, Rashna Bhandari

## Abstract

The inositol pyrophosphate 5-InsP_7_, composed of an inositol ring substituted with five monophosphates and one diphosphate, modulates diverse cellular functions by protein pyrophosphorylation, during which its β-phosphate moiety is transferred to a pre-phosphorylated serine residue on the target protein. In mammals, the synthesis of 5-InsP_7_ from its precursor InsP_6_ is catalyzed by a family of enzymes called IP6Ks. We report that during recovery from genotoxic stress, cells lacking the IP6K isoform IP6K1 exhibit prolonged persistence of DNA damage foci marked by the homologous recombination repair protein RAD51. Expression of catalytically active but not inactive IP6K1 reverses this defect, implying that 5-InsP_7_ supports the dissolution of RAD51 foci. Upon DNA damage, we observe an increase in IP6K1 activity, contingent on its phosphorylation by the protein kinases CK2 and CDK1. IP6K1 is recruited to sites of DNA damage, and interacts with RAD51, CDK1, and the C-terminal domain (CTD) of BRCA2. Disruption of binding between RAD51 and BRCA2-CTD is known to support the disassembly of RAD51 foci. We show that 5-InsP_7_ can pyrophosphorylate RAD51, and that the presence of 5-InsP_7_ diminishes RAD51 binding to BRCA2-CTD. Our findings provide a mechanism by which 5-InsP_7_ synthesized by IP6K1 facilitates the removal of RAD51 from sites of DNA repair.

**Summary statement:** Inositol hexakisphosphate kinase 1, an enzyme that catalyses the synthesis of the inositol pyrophosphate 5-InsP_7_, localises to DNA double strand breaks, and engages in interactions with proteins involved in homologous recombination (HR)-mediated DNA repair. 5-InsP_7_ disrupts the interaction between RAD51 and the C-terminus of BRCA2, promoting dislodgement of RAD51 from DNA damage sites post-repair.

## Introduction

Inositol pyrophosphates are a class of energy-rich small signalling molecules that are found ubiquitously in all eukaryotes. Inositol hexakisphosphate (InsP_6_) kinases (IP6Ks) catalyse the synthesis of the inositol pyrophosphate 5-PP-InsP_5_ (5-InsP_7_) from InsP_6_ [1–4]. Mammals have three IP6K paralogs named IP6K1/K2/K3, whereas KCS1 is the only InsP_6_ kinase in *S. cerevisiae* [1–6]. Inositol pyrophosphates are known to modulate several cellular processes, primarily via two molecular mechanisms - (i) regulation of proteins by direct binding, or (ii) by orchestrating a non-canonical post-translational modification called protein pyrophosphorylation [1, 2, 7–11]. During the latter process, 5-InsP_7_ transfers its β-phosphate moiety to a pre-phosphorylated Ser residue that lies in the vicinity of Glu/Asp residues within an intrinsically disordered region of the target protein [7, 8, 11–18]. The priming phosphorylation is usually brought about by acidophilic serine / threonine kinases such as CK2 [7, 8, 17, 18]. Pyrophosphorylation of a protein by 5-InsP_7_ is an enzyme-independent modification, but requires the presence of Mg^2+^ ions, and is assisted by interaction of the target protein with an IP6K, which increases the local concentration of 5-InsP_7_ to drive the reaction [8, 18].

5-InsP_7_ plays a pivotal role in mammalian and yeast physiology. It regulates various metabolic and signalling pathways, including rRNA synthesis, vesicle trafficking, immune signalling, apoptosis, and insulin exocytosis [1, 2, 8, 9, 11, 14, 17, 19]. InsP_6_ kinases are reported to be involved in the maintenance of genome integrity in budding yeast and mammalian cells [20–23]. 5-InsP_7_ synthesized by the yeast InsP_6_ kinase KCS1 has been shown to support DNA recombination [24, 25]. In mammals, 5-InsP_7_ synthesized by IP6K1 upregulates both the nucleotide excision repair and homologous recombination (HR) repair pathways [22, 23]. In earlier work, we have shown that in comparison with their wild type counterparts, mouse embryonic fibroblasts (MEFs) lacking *Ip6k1* are able to initiate DNA repair when treated with the genotoxic agents hydroxyurea (HU) or neocarzinostatin (NCS), but exhibit reduced viability post drug removal [22]. Additionally, *Ip6k1^-/-^* MEFs display persistent DNA damage foci marked by phosphorylated histone H2AX (γH2AX), and the HR repair proteins BLM and RAD51, even after removal of the DNA damage agent. This function of IP6K1 in HR-mediated DNA repair is dependent on its ability to synthesize 5-InsP_7_, but the molecular mechanism by which this inositol pyrophosphate facilitates removal of repair proteins from DNA damage sites remains elusive.

In the present study, we have investigated the molecular mechanism underlying the influence of IP6K1 on dissolution of DNA repair foci. We show that cells lacking IP6K1 exhibit persistent RAD51 foci during recovery from DNA damage induced by the inter-strand crosslinker mitomycin C (MMC). DNA damage induces an increase in IP6K1 activity in response to phosphorylation catalysed by protein kinases CK2 and CDK1-CCNB1 (cyclin B1). We show that IP6K1 is recruited near the site of DNA damage, and interacts with the DNA repair proteins RAD51, CDK1, and BRCA2. 5-InsP_7_ synthesized by IP6K1 pyrophosphorylates RAD51 in its N-terminal disordered region, disrupting RAD51 binding with the C-terminal region of BRCA2, and supporting the removal of RAD51 foci post DNA repair.

## Results

### Depletion of IP6K1 leads to persistence of DNA damage foci

We have previously shown that recovery from DNA damage via the HR pathway is stalled in MEFs lacking IP6K1 [22]. Upon prolonged treatment with the replication stressor HU, HR repair proteins were recruited to nuclear foci in *Ip6k1*^-/-^ MEFs, but following removal of the drug, the damage foci persisted in *Ip6k1*^-/-^ MEFs although they were cleared in *Ip6k1*^+/+^ MEFs. In the present study, to uncover the molecular basis underlying IP6K1 function in HR repair, we used the human osteosarcoma cell line U-2 OS, a p53 harbouring cell line that has been used widely to study DNA repair pathways [26, 27]. U-2 OS cells expressing shRNA directed against *IP6K1* (sh*IP6K1*) exhibit ∼65% knockdown in IP6K1 expression compared with control cells that harbour a non-targeting shRNA (shNT) (Supplementary Figure S1A) [28]. We monitored the effect of IP6K1 depletion on InsP_7_ levels by employing two different methods to measure the cellular content of InsP_6_ and InsP_7_ (Supplementary Figure S1B - D). In the first method, the total cellular pools of InsP_6_ and InsP_7_ were enriched on titanium dioxide (TiO_2_) beads and visualised by PAGE analysis [29]. In the second method, we utilised [^3^H]-inositol to label the pool of inositol phosphates and resolved these by strong anion exchange HPLC [30]. Both analyses revealed a decrease in InsP_7_ in U-2 OS sh*IP6K1* cells compared with shNT cells, with no significant change in the level of InsP_6_. [^3^H]-inositol labelling confirmed that depletion of IP6K1 did not affect the levels of phosphatidylinositol (PtdIns) lipids (Supplementary Figure S1D).

To study the role of IP6K1 in DNA repair, we subjected U-2 OS shNT and sh*IP6K1* cells to treatment with different genotoxic stressors that are known to trigger the HR repair pathway [31]. Cells treated with the drug were allowed to recover, and DNA damage foci were scored by immunostaining for γH2AX, a damage-specific marker [32]. Following treatment with the inter-strand crosslinker mitomycin C (MMC) [33], we observed the accumulation of equivalent γH2AX foci in U-2 OS shNT and sh*IP6K1* cells, but noticed a stark difference in the foci count during recovery from the drug (Figure 1A and B). Following drug removal, shNT cells showed a substantial decrease in the number of damage foci after 12 hours, whereas sh*IP6K1* cells, while showing some recovery, exhibited a much higher number of persistent DNA damage foci in their nuclei. Cells treated with the radiomimetic drug NCS [34] or the DNA intercalating agent, phleomycin [35] exhibited the same pattern. In both cases, U-2 OS shNT cells recovered significantly from DNA damage within 8 h after removal of the drug, but sh*IP6K1* cells showed a much lower extent of recovery at this time (Figure 1C - F). The inability of cells lacking IP6K1 to recover from DNA damage induced by three different agents highlights the pivotal role exerted by IP6K1 in DNA repair.

**Figure 1.**
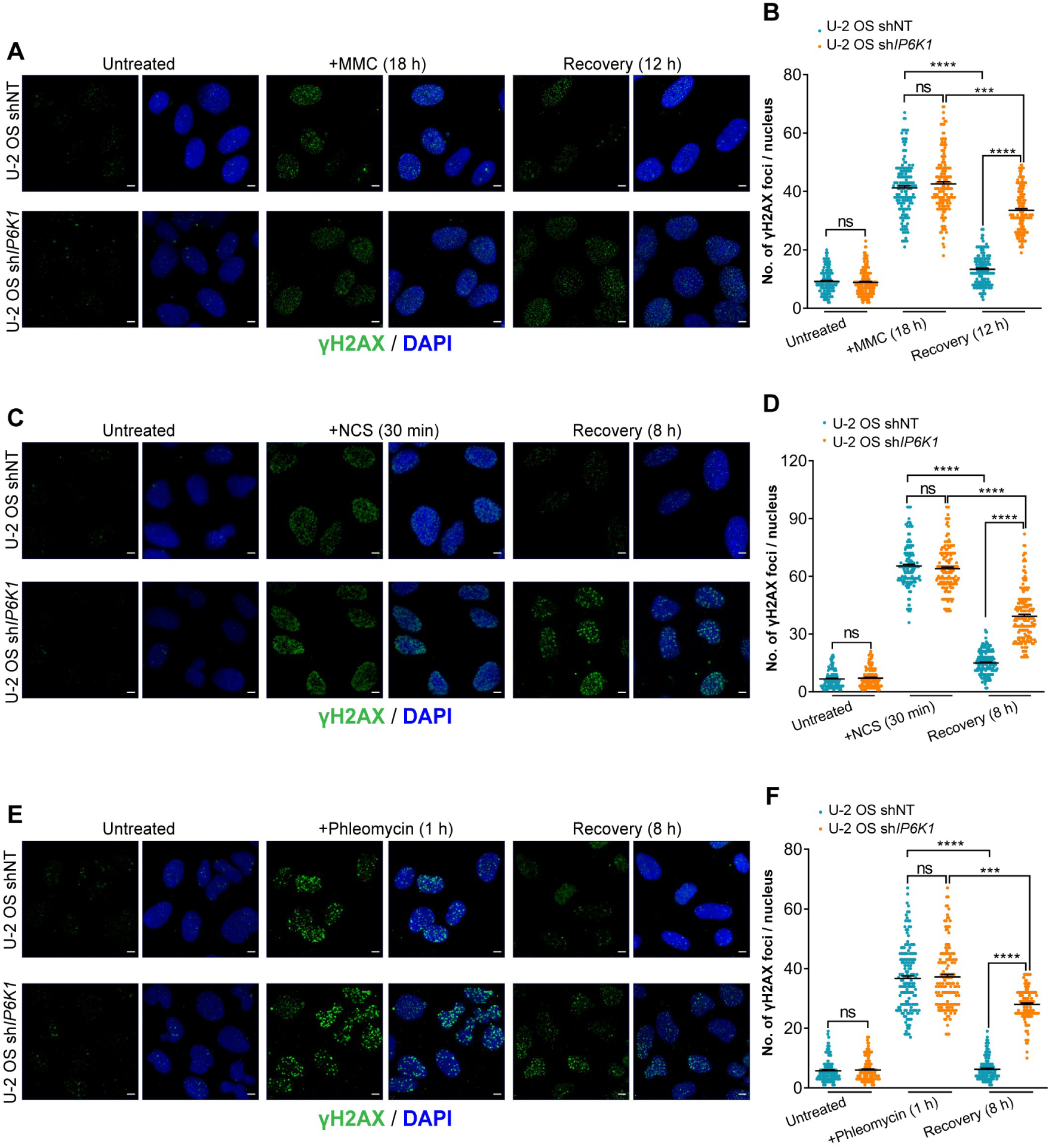
DNA damage persists in cells depleted for IP6K1. (**A**) Representative immunofluorescence images of U-2 OS shNT and sh*IP6K1* cells, left untreated, treated with mitomycin C (MMC; 0.5µM), or allowed to recover from MMC-induced DNA damage for the indicated time (post-recovery), and stained with anti-γH2AX antibody (green). **(B)** Quantification of the number of γH2AX foci per nucleus in (A). Data distribution is represented by scatter dot plots (*n* = 155, 167, and 174 nuclei for untreated, MMC treated, and post-recovery U-2 OS shNT cells, and *n* = 163, 172, and 172 nuclei for untreated, MMC treated, and post-recovery U-2 OS sh*IP6K1* cells, from three independent experiments). **(C)** Representative immunofluorescence images of U-2 OS shNT and sh*IP6K1* cells, left untreated, treated with neocarzinostatin (NCS; 200 µg/mL), or allowed to recover from NCS-induced DNA damage for the indicated time, and stained with anti-γH2AX antibody (green). **(D)** Quantification of the number of γH2AX foci per nucleus in (C). Data distribution is represented by scatter dot plots (*n* = 146, 171, and 181 for untreated, NCS treated, and post-recovery U-2 OS shNT cells, and *n* = 172, 168, and 181 for untreated, NCS treated and post-recovery U-2 OS sh*IP6K1* cells, from three independent experiments). **(E)** Representative immunofluorescence images of U-2 OS shNT and sh*IP6K1* cells, left untreated, treated with phleomycin (10 µM), or allowed to recover from phleomycin-induced DNA damage for the indicated time, and stained with anti-γH2AX antibody (green). **(F)** Quantification of the number of γH2AX foci per nucleus in (E). Data distribution is represented by scatter dot plots (*n* = 147, 166, and 138 for untreated, phleomycin treated, and post-recovery U-2 OS shNT cells, and *n* = 128, 149, and 144 for untreated, phleomycin treated, and post-recovery U-2 OS sh*IP6K1* cells, from three independent experiments). Nuclei were stained with DAPI (blue) and scale bars are 5 μm in A, C, and E. Data presented in B, D, and F were analyzed using Kruskal-Wallis test with Dunn’s multiple comparison post hoc test; ns, non-significant (*P* > 0.05), \*\*\**P* ≤ 0.001, \*\*\*\**P* ≤ 0.0001.

### Disassembly of DNA damage-induced RAD51 foci requires 5-InsP_7_ synthesis by IP6Ks

HR-mediated DNA repair is a complex multistep process which involves the stage-wise recruitment and removal of several proteins [36]. RAD51, a central player in HR, is recruited to the sites of DNA damage in a BRCA2 dependent manner [37, 38]. RAD51 forms a nucleoprotein filament on the overhanging single stranded DNA (ssDNA), searches for the homologous template strand, and participates in strand invasion. We counted nuclear RAD51 foci as a surrogate marker to study the course of HR-mediated repair in U-2 OS cells treated with MMC. RAD51 foci accumulation following MMC treatment was comparable in shNT and sh*IP6K1* U-2 OS cells, but post drug removal, IP6K1 depleted cells displayed persistent RAD51, whereas shNT cells exhibited a significant reduction in RAD51 foci (Figure 2A and B). As sh*IP6K1* U-2 OS cells still express low levels of IP6K1, we employed CRISPR-Cas9 technology to generate *IP6K1* knockout (KO) U-2 OS cell lines. Two independent *IP6K1* KO U-2 OS cell lines were obtained, both of which showed significantly lower levels of InsP_7_ compared with non-targeted control (NTC) U-2 OS cells (Supplementary Figure S1E - J). As seen with the sh*IP6K1* U-2 OS cell line, both *IP6K1* KO cell lines showed persistent RAD51 foci during recovery from MMC treatment (Supplementary Figure S2A and B). To evaluate whether impaired RAD51 foci disassembly in the absence of IP6K1 is conserved across cell lines, we generated two independent *IP6K1* KO HEK293T cell lines (Supplementary Figure S1K; [17]. Compared with wild type HEK293T cells both *IP6K1* KO lines displayed a substantial reduction in InsP_7_ (Supplementary Figure S1L - N; [17]. The phenomenon of RAD51 foci persistence during recovery from MMC treatment was conserved in both *IP6K1* KO HEK293T cell lines (Supplementary Figure S2C and D).

**Figure 2.**
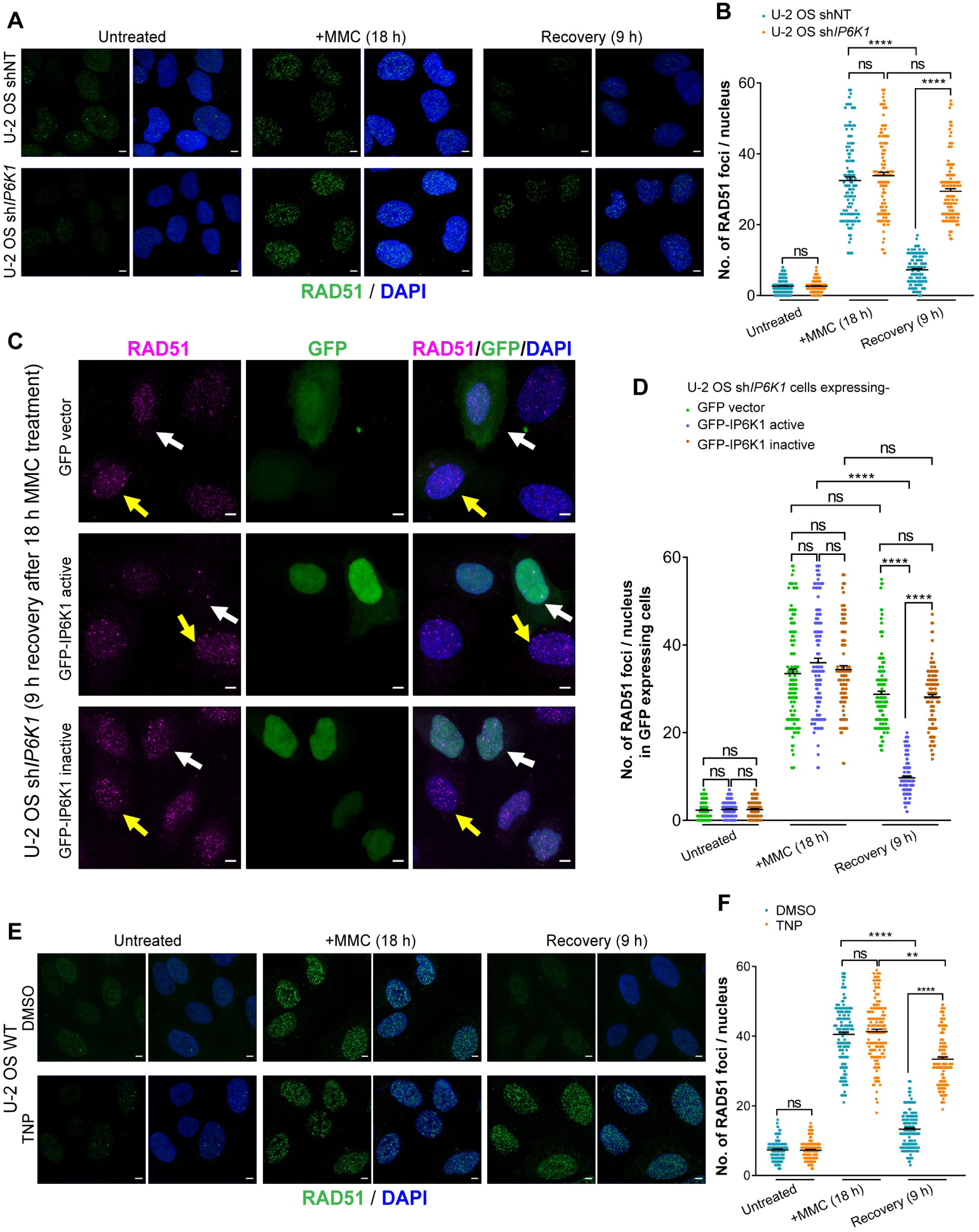
5-InsP_7_ promotes RAD51 foci dissolution. **(A)** Representative immunofluorescence images of U-2 OS shNT and sh*IP6K1* cells, left untreated, treated with MMC, or allowed to recover from MMC-induced DNA damage, and stained with anti-RAD51 antibody (green). **(B)** Quantification of the number of RAD51 foci per nucleus in (A). Data distribution is represented by scatter dot plots (*n* = 132, 125, and 128 nuclei for untreated, MMC treated, and post-recovery U-2 OS shNT cells, and *n* = 143, 138, and 135 nuclei for untreated, MMC treated, and post-recovery U-2 OS sh*IP6K1* cells, from four independent experiments). **(C)** Representative immunofluorescence images of U-2 OS sh*IP6K1* cells overexpressing GFP (green), GFP-tagged active mouse IP6K1, or inactive (K226A/S334A) mouse IP6K1, treated with MMC and allowed to recover for 9 h, stained with anti-RAD51 antibody (magenta). Transfected and untransfected cells are indicated with white and yellow arrows respectively. Representative immunofluorescence images corresponding to untreated and MMC treated cells are presented in Supplementary Figure S2E and F. **(D)** Quantification of the number of RAD51 foci per nucleus in (C), S2E and S2F. Data distribution is represented by scatter dot plots (*n* = 126, 132, and 118 nuclei for untreated U-2 OS sh*IP6K1* cells expressing vector, IP6K1 (active), and IP6K1 (inactive); *n* = 125, 140, and 122 nuclei for MMC treated U-2 OS sh*IP6K1* cells expressing vector, IP6K1 (active), and IP6K1 (inactive); and *n* = 135, 139, and 125 nuclei for post-recovery U-2 OS sh*IP6K1* cells expressing vector, IP6K1 (active), and IP6K1 (inactive), from four independent experiments). **(E)** Representative immunofluorescence images of U-2 OS wild-type (WT) cells pre-treated with DMSO (vehicle control) or TNP (10 µM for 16 h), followed by no DNA damage (untreated), MMC treatment, or recovery from MMC-induced DNA damage, in the continued presence of DMSO or TNP as indicated, and stained with anti-RAD51 antibody (green). **(F)** Quantification of the number of RAD51 foci per nucleus in (E). Data distribution is represented by scatter dot plots (*n* = 148, 152, and 170 for untreated, MMC treated and post-recovery U-2 OS WT in DMSO; and *n* = 163, 159, and 158 for untreated, MMC treated and post-recovery U-2 OS WT cells in TNP, from three independent experiments). Nuclei were stained with DAPI (blue), and scale bars are 5 μm in A, C, and E. Data in B, D, and F were analyzed using Kruskal-Wallis test with Dunn’s multiple comparison post hoc test; ns, non-significant, (*P* > 0.05), \*\**P* ≤ 0.01, \*\*\*\**P* ≤ 0.0001.

To determine whether the catalytic activity of IP6K1 is essential for the disassembly of RAD51 foci, we ectopically introduced GFP-tagged active or inactive (K226A/S334A) IP6K1 into U-2 OS sh*IP6K1* cells (Supplementary Figure S3A - C). No difference was observed in the number of RAD51 foci in untreated or MMC treated U-2 OS sh*IP6K1* cells expressing active or inactive IP6K1 (Supplementary Figure S2E and F). However, following the removal of MMC, cells expressing catalytically proficient IP6K1 showed a significant reduction in RAD51 foci, whereas cells expressing inactive IP6K1 showed no recovery (Figure 2C and D). To lend further support to the requirement of IP6K enzyme activity for RAD51 foci disassembly, we employed the pan-IP6K pharmacological inhibitor, TNP [39] (Supplementary Figure S3D and E). Upon removal of MMC, vehicle treated control cells displayed a significant reduction in RAD51 foci in 9 hours, whereas in TNP treated cells there was a much higher number of persistent RAD51 foci (Figure 2E and F). These findings highlight the essentiality of 5-InsP_7_ for RAD51 foci disassembly during recovery from MMC treatment.

### IP6K1 activity increases upon DNA damage and recovery

To examine how DNA damage signalling influences IP6K1, we monitored the localization, level and activity of IP6K1 following MMC treatment and recovery. We have earlier shown that in U-2 OS cells, IP6K1 is detected in the cytoplasm and nucleus, and is especially enriched in the fibrillar center of nucleoli [17, 28]. Proteins involved in DNA damage repair are known to localise at the sites of DNA damage when cells are exposed to genotoxic stressors [40]. To investigate whether IP6K1 shows this behaviour we co-stained U-2 OS shNT cells to detect γH2AX and IP6K1. As reported earlier [17], IP6K1 was seen throughout the cell with intense staining in nuclear foci (Figure 3A and B). There was no apparent change in IP6K1 localisation upon MMC treatment or recovery, and no evidence of co-localisation of IP6K1 with γH2AX foci. Additionally, we investigated whether DNA damage and recovery influences the cellular levels of IP6K1. We confirmed that γH2AX levels increased upon MMC treatment and decreased during recovery, reflecting activation of the DNA damage signalling cascade, but noted that IP6K1 levels remained unaltered across these time points (Figure 3C and D). Next, we tested whether MMC-induced DNA damage has an effect on IP6K1 enzyme activity. We first demonstrated that IP6K1 immunoprecipitated from HEK293T cells is able to synthesize 5-InsP_7_ from InsP_6_ and ATP (Supplementary Figure S3F). The activity of endogenous IP6K1 increased ∼1.5 fold following MMC treatment and remained high during the recovery phase (Figure 3E and F). We then explored the possibility that changes in phosphorylation may be a mechanism by which IP6K1 activity is upregulated following MMC treatment. We focused on the protein kinases CK2, Polo-like kinase 1 (PLK1), and CDK1-Cyclin B1 (CCNB1), as the former two kinases are known to regulate RAD51 foci formation on sites of DNA damage [41], and the latter is known to participate in the disassembly of RAD51 nucleoprotein filaments [42]. All three kinases catalysed *in vitro* phosphorylation of GST-tagged IP6K1 expressed in *E. coli*, whereas GST alone did not undergo phosphorylation (Figure 3G). Phosphorylation of IP6K1 by CK2 or CDK1-CCNB1 resulted in elevation of 5-InsP_7_ synthesis compared with unphosphorylated IP6K1, whereas PLK1 phosphorylation did not affect IP6K1 activity (Figure 3H and I). These data suggest that a phosphorylation-dependent increase in IP6K1 activity may augment 5-InsP_7_ availability to support disassembly of damage-induced RAD51 foci.

**Figure 3.**
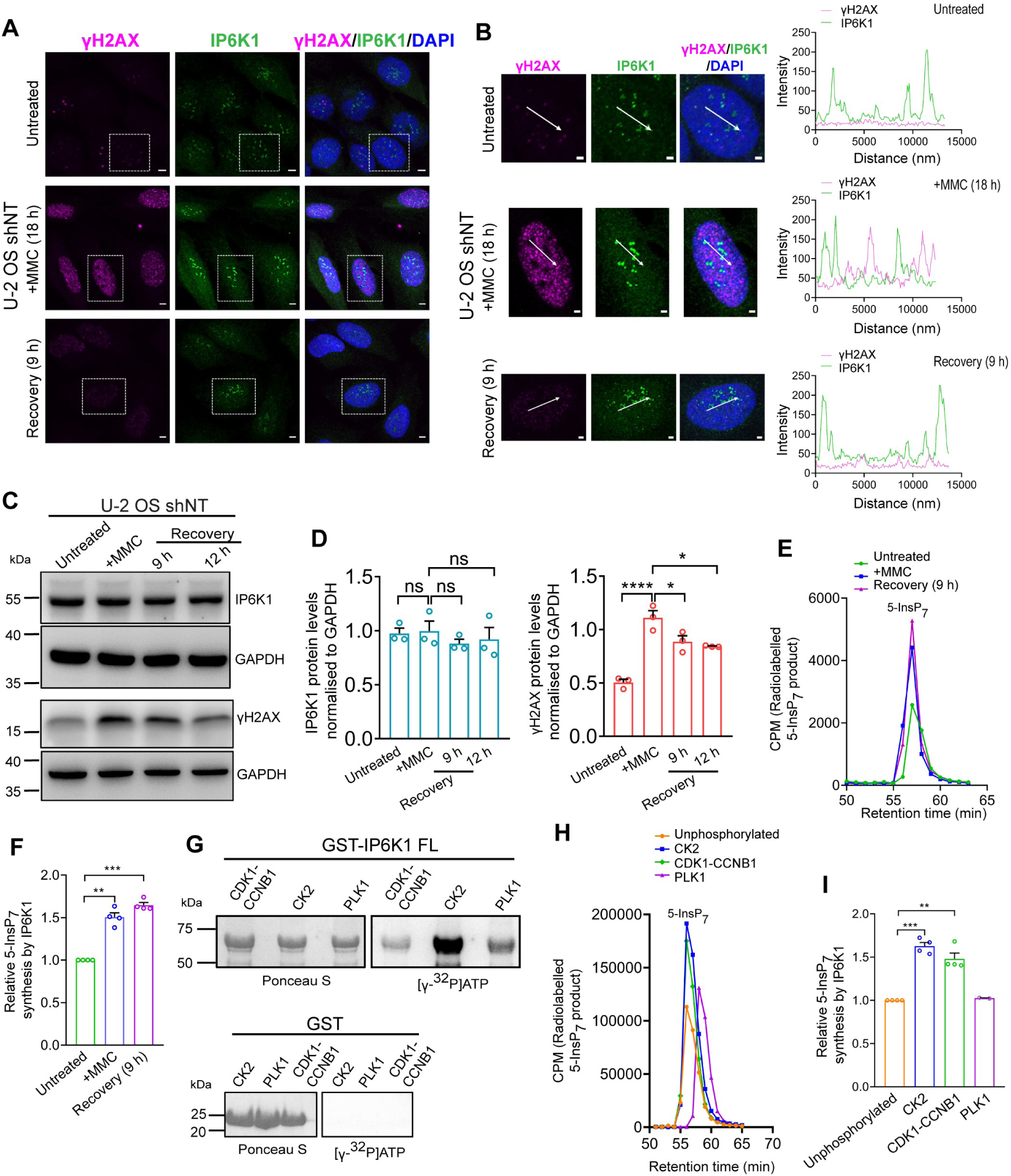
Effect of DNA damage on IP6K1. **(A)** Localisation of endogenous IP6K1 (green) and γH2AX (magenta) in U-2 OS shNT cells, left untreated, treated with MMC, or allowed to recover from MMC-induced DNA damage. Nuclei were stained with DAPI (blue). **(B)** The boxed regions in (A) were magnified and fluorescence intensity profiles for IP6K1 (green) and γH2AX (magenta) were measured along the indicated white arrows. Scale bars are 5 μm in A and 2 μm in B. Co-localization of γH2AX with IP6K1 was assessed by calculating Pearsons’s correlation coefficient, which yielded values of 0.128 ± 0.019, 0.150 ± 0.018, and 0.132 ± 0.013 (mean ± S.E.M., *n* = 50 in each case) for untreated, MMC treated, and post-recovery cells, respectively, indicating no significant co-localization. **(C)** Representative immunoblots showing IP6K1 and γH2AX levels in U-2 OS shNT cells upon MMC treatment and the indicated time following MMC removal. GAPDH was used as a loading control. **(D)** Quantification of (C). Data (mean ± SEM, *N* = 3), depicting IP6K1 and γH2AX protein levels normalised to GAPDH, were analyzed using ordinary one-way ANOVA with Tukey’s multiple comparisons test; ns, non-significant (*P* > 0.05), \**P* ≤ 0.05, \*\*\*\**P* ≤ 0.0001. **(E)** Chromatogram traces depicting the SAX-HPLC analysis of 5[β-^32^P]InsP_7_ formed during incubation of endogenous IP6K1 immunoprecipitated from untreated, MMC treated, or post-recovery HEK293T cells with [γ-^32^P]ATP and InsP_6._ **(F)** Quantification of (E). The extent of 5[β-^32^P]InsP_7_ synthesis by IP6K1 from MMC treated and post-recovery cells, normalised to untreated cells. Data (mean ± SEM, *N* = 4), were analyzed using a one-sample *t* test, ** *P* ≤ 0.01, \*\*\**P* ≤ 0.001. **(G)** GST and GST-tagged full length (FL) human IP6K1 expressed in *E. coli* were incubated with CDK1-CCNB1, CK2, or PLK1 in presence of [γ-^32^P]ATP. Representative images show autoradiography to detect phosphorylation mediated by protein kinases (right) and GST-tagged proteins were detected by Ponceau S staining (left) (*N* = 3). **(H)** Chromatogram traces depicting SAX-HPLC analysis of 5[β-^32^P]InsP_7_ formed during incubation of GST-tagged human IP6K1 (GST-IP6K1) with [γ-^32^P]ATP and InsP_6_, subsequent to phosphorylation by CDK-CCNB1, CK2, or PLK1. **(I)** Quantification of (H). The extent of 5[β-^32^P]InsP_7_ synthesis by GST-IP6K1 following phosphorylation with the abovementioned kinases, normalised to the activity of unphosphorylated GST-IP6K1. Data (mean ± SEM, *N* = 3 for CDK1-CCNB1 and CK2, and N = 2 for PLK1), were analyzed using a one-sample *t* test, ** *P* ≤ 0.01, \*\*\**P* ≤ 0.001.

### Phosphorylation of IP6K1 within a disordered region in the N-lobe elevates its activity

To identify the region of IP6K1 that undergoes phosphorylation by CK2 and CDK1-CCNB1, we generated GST-fusions of different fragments of IP6K1 based on disorder prediction by the IUPred3 tool (https://iupred3.elte.hu/) and tertiary structure prediction using AlphaFold Monomer v2.0 (https://alphafold.ebi.ac.uk/entry/AF-Q92551-F1) (Figure S3G). Our expression constructs correspond to the amino (N-) terminal or carboxyl (C-) terminal lobes of IP6K1, and intrinsically disordered regions, designated IDR1 and IDR2, that lie within the N-and C-lobes respectively (Supplementary Figures S3G and S4A). *In vitro* phosphorylation of these GST-tagged IP6K1 fragments revealed that both CK2 and CDK1-CCNB1 phosphorylate residues lying within IDR1 in the N-lobe (Figure 4A and B). Next, we employed mass spectrometry to identify phosphosites on IP6K1 isolated from HEK293T cells under three different conditions – cells that are untreated, treated with MMC, or allowed to recover following MMC treatment. Seven phosphosites were identified on IP6K1 across these conditions, of which six sites (Ser125, Ser127, Ser129, Ser139, Thr142, and Ser146) lie within IDR1, and one site (Ser381) is within IDR2 (Supplementary Figure S4B and C). Phosphorylation on these sites under different treatment conditions was detected with varying statistical confidence, with only Thr142 phosphorylation seemingly dependent on MMC treatment (Supplementary Figure S4D). To determine whether any of these sites are the targets of CK2 or CDK1-CCNB1, we replaced each of these seven Ser/Thr residues individually with Ala and subjected the mutant IP6K1 proteins to phosphorylation by CK2 or CDK1-CCNB1. We observed no significant change in the extent of phosphorylation of these point mutant forms of IP6K1 compared with the native sequence (Supplementary Figure S5A and B). This suggests that the principal target site for these kinases may be different from the phosphosites identified by mass spectrometry, or that these protein kinases may phosphorylate multiple sites within IDR1, which contains 18 Ser/Thr residues (Figure S5D).

**Figure 4.**
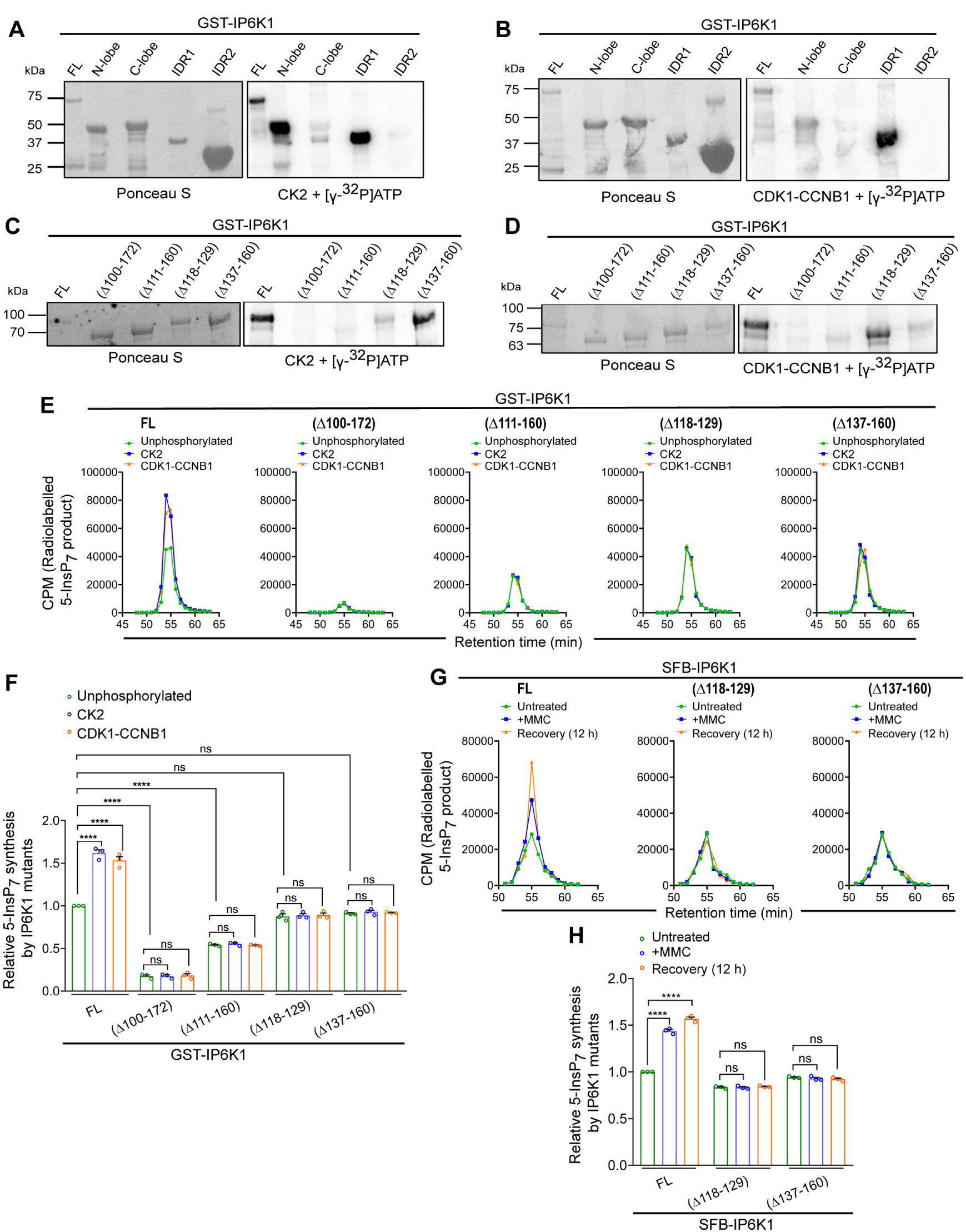
N-lobe phosphorylation enhances IP6K1 activity. **(A-D)** GST-IP6K1 domains (A, B; see Figure S4A) and in-frame IDR1 deletion mutants (C, D; see Figure S5C) expressed in *E. coli* were subjected to phosphorylation by either CK2 (A, C) or CDK1-CCNB1 (B, D) in presence of [γ-^32^P]ATP. Representative images show autoradiography to detect phosphorylation mediated by protein kinases (right) and GST-tagged proteins stained with Ponceau S (left) (*N* = 2). **(E)** Chromatogram traces depicting SAX-HPLC analysis of 5[β-^32^P]InsP_7_ formed during incubation of IDR1 deletion mutants of GST-IP6K1 with [γ-^32^P]ATP and InsP_6_, subsequent to phosphorylation by CK2 or CDK1-CCNB1. **(F)** Quantification of (E). The extent of 5[β-^32^P]InsP_7_ synthesis by GST-IP6K1 mutants following phosphorylation with the abovementioned kinases, normalised to the activity unphosphorylated full length (FL) GST-IP6K1. Data (mean ± SEM, *N* = 3), were analyzed using ordinary one-way ANOVA with Tukey’s multiple comparisons test; ns, non-significant (*P* > 0.05), \**P* ≤ 0.05, \*\*\*\**P* ≤ 0.0001. **(G)** Chromatogram traces depicting the SAX-HPLC analysis of 5[β-^32^P]InsP_7_ formed during incubation of IDR1 deletion mutants of SFB-tagged human IP6K1 expressed in HEK293T cells with [γ-^32^P]ATP and InsP_6_, subsequent to MMC treatment and post-recovery. **(H)** Quantification of (G). The extent of 5[β-^32^P]InsP_7_ synthesis by SFB-IP6K1 mutants following MMC treatment and post-recovery, normalised to the activity of untreated full length (FL) SFB-IP6K1. Data (mean ± SEM, *N* = 3), were analyzed using ordinary one-way ANOVA with Tukey’s multiple comparisons test; ns, non-significant (*P* > 0.05), \*\*\*\**P* ≤ 0.0001.

Therefore, we adopted a different approach to narrow down the region in IP6K1 that undergoes phosphorylation by CK2 and CDK1-CCNB1. We tested phosphorylation of in-frame deletion mutants of IP6K1 in which different segments of IDR1 were removed (Supplementary Figure S5C and D). In the IP6K1(Δ100-172) mutant in which IDR1 has been completely truncated, phosphorylation by both CK2 and CDK1-CCNB1 was eliminated, confirming that all phosphosites for these protein kinases lie within IDR1 (Figure 4C and D). Notably, removal of IDR1 nearly abolished IP6K1 enzyme activity (Figure 4E and F). As expected, incubation with CK2 or CDK1-CCNB1 had no effect on the activity of the IP6K1(Δ100-172) mutant. A shorter in-frame deletion of IDR1(Δ111-160) also abrogated phosphorylation by CK2 and CDK1-CCNB1 (Figure 4C and D). Although InsP_6_ kinase activity in this mutant was only reduced by approximately 50% compared with the native protein, pre-incubation with either CK2 or CDK1-CCNB1 had no effect on 5-InsP_7_ synthesis (Figure 4E and F). Residues 111-160 of IP6K1 contain two clusters of potential phosphosites – one cluster from Ser118 to Ser129, and another cluster from Ser137 to Ser160 (Supplementary Figure S5D). We generated independent in-frame deletion mutants for these Ser clusters, IP6K1(Δ118-129) and IP6K1(Δ137-160), and tested both mutant constructs for phosphorylation by CK2 and CDK1-CCNB1. The IP6K1(Δ118-129) mutant showed significantly lower phosphorylation by CK2 compared with full length IP6K1, whereas the IP6K1(Δ137-160) mutant showed a similar extent of phosphorylation as the native sequence (Figure 4C). Conversely, phosphorylation by CDK1-CCNB1 was substantially reduced in the IP6K1(Δ137-160) mutant, but was preserved in the IP6K1(Δ118-129) mutant (Figure 4D). These results clearly demonstrate that CK2 and CDK1-CCNB1 phosphorylate independent regions within IDR1, which lie between Ser118-Ser129 and Ser137-Ser160 respectively. Finally, we tested the effect of phosphorylation by CK2 or CDK1-CCNB1 on the catalytic activity of both Ser cluster IP6K1 mutants. The (Δ118-129) and (Δ137-160) mutant forms of IP6K1 retain nearly the same activity as the native protein (Figure 4E and F). As expected, incubation with CK2 did not enhance the 5-InsP_7_ synthesis activity of IP6K1(Δ118-129). Interestingly, despite phosphorylating IP6K1(Δ118-129), CDK1-CCNB1 failed to upregulate its catalytic activity. Similarly, the catalytic activity of IP6K1(Δ137-160) was unaffected by either CK2 or CDK1-CCNB1. These data show that although CK2 and CDK1-CCNB1 phosphorylate independent clusters within IDR1, the entire IDR1 sequence is necessary for upregulation of IP6K1 activity in response to phosphorylation by these protein kinases.

Next, we introduced the in-frame deletions (Δ118-129) and (Δ137-160) into SFB-tagged IP6K1 for expression in mammalian cells. As seen in the case of endogenous IP6K1 (Figure 3E and F), SFB-IP6K1 expressed in HEK293T cells showed increased catalytic activity following MMC treatment and recovery, compared with IP6K1 isolated from untreated cells (Figure 4G and H). However, IP6K1(Δ118-129) and IP6K1(Δ137-160) did not show any change in 5-InsP_7_ synthesis activity in response to MMC treatment and recovery. These data suggest that CK2 and CDK1-CCNB1 phosphorylation may be responsible for upregulating IP6K1 activity in response to MMC-induced DNA damage.

### IP6K1 is recruited to DNA damage sites and interacts with repair proteins

To explore the mechanism by which IP6K1 induces the disassembly of DNA damage foci, we first checked if IP6K1 affects the efficiency of HR-mediated repair by employing a reporter assay. We used a U-2 OS cell line that carries two mutated copies of GFP oriented as direct repeats (DR-GFP; [27, 43]; Supplementary Figure S6A). The 5’ GFP sequence harbors a cleavage site for the endonuclease I-SceI and two premature stop codons (SceGFP), and the 3’ GFP sequence contains upstream and downstream truncations that render it inactive (iGFP). These cells also express I-SceI fused with the glucocorticoid receptor ligand binding domain and mCherry tag (mCherry-I-SceI-GR), which is translocated to the nucleus upon treatment of cells with the glucocorticoid receptor agonist triamcinolone acetonide. An I-SceI-mediated double strand break in SceGFP is repaired by HR using iGFP as a template, generating a sequence encoding active GFP, which can be assessed by flow cytometry. To our surprise, the depletion of *IP6K1* in U-2 OS DR-GFP cells did not lead to a reduction in GFP expression compared with non-targeted U-2 OS DR-GFP cells, suggesting that IP6K1 does not directly regulate HR efficiency (Figure 5A and B). As a positive control, we confirmed that *RAD51* depletion in these cells leads to a reduction in GFP expression reflecting an impairment in HR. Next, we employed the U-2 OS DR-GFP cell line to monitor the recruitment of IP6K1 to I-SceI-generated DNA breaks, by conducting a chromatin immunoprecipitation (ChIP) on break assay [44]. We examined the binding of IP6K1 and RAD51 to three regions of DNA at different distances from the I-SceI cut site (Supplementary Figure S6B). As shown in earlier studies [44, 45], RAD51 was enriched proximal to the DNA break at ∼100 bp from the cut site, but showed reduced abundance at more distal sites centered at ∼250 bp and ∼1.1 kb from the cut site (Figure 5C and Supplementary Figure S6B). Upon induction of DNA breaks, IP6K1 showed enhanced occupancy on DNA, with maximum association at ∼250 bp from the cut site (Figure 5C). These data suggest that IP6K1 does not directly regulate HR efficiency but binds near DNA damage sites and may influence RAD51 foci disassembly by locally increasing the level of 5-InsP_7_.

**Figure 5.**
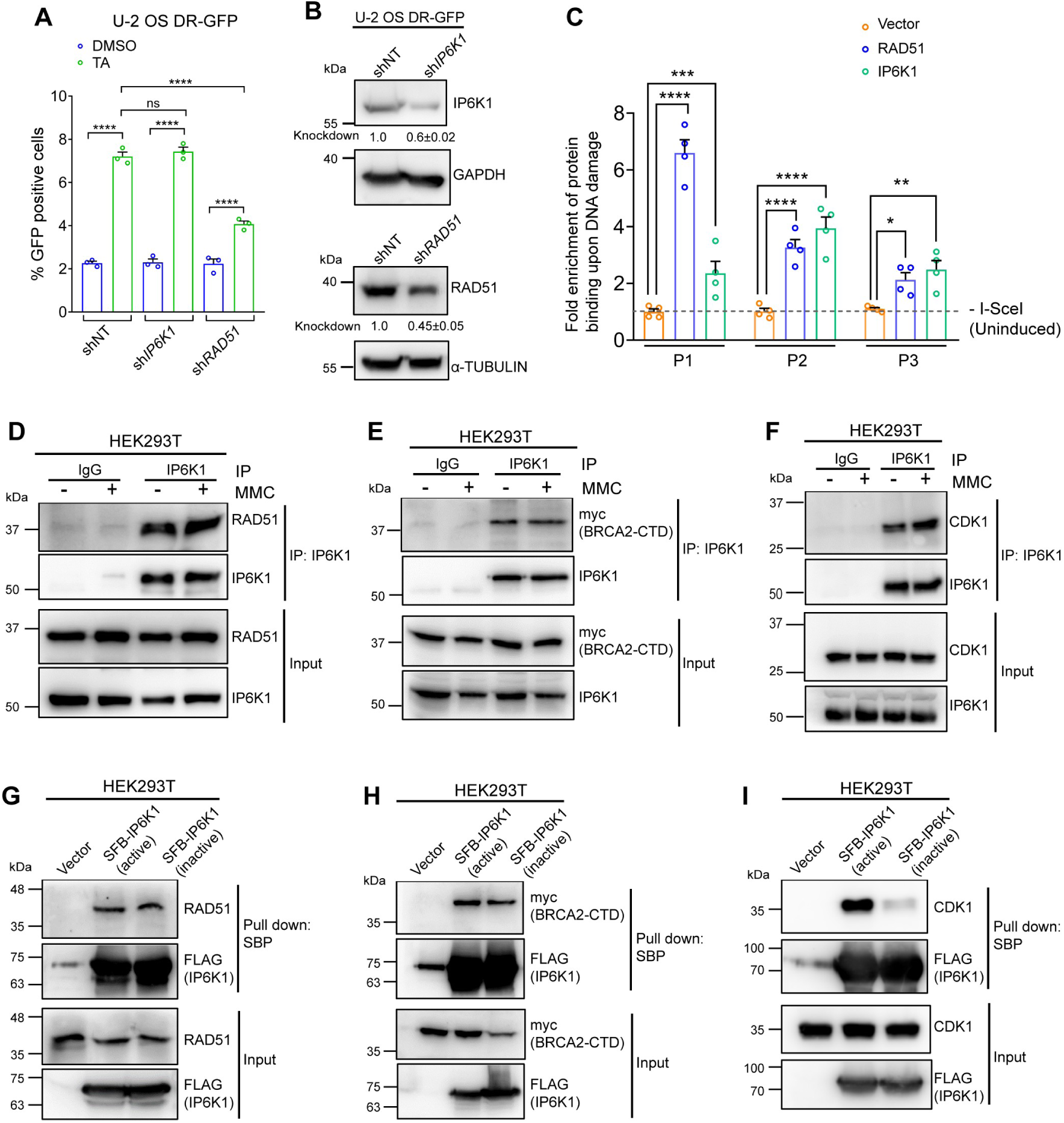
IP6K1 interacts with DNA repair proteins. **(A)** Flow cytometry analysis of U-2 OS DR-GFP cells expressing non-targeting control shRNA (shNT), or shRNA against human *IP6K1* (sh*IP6K1*) or *RAD51* (sh*RAD51*), following treatment with DMSO (vehicle control) or triamcinolone acetonide (TA) to induce expression of I-SceI (see Supplementary Figure S6A). Data (mean ± SEM, *N* = 3), were analyzed using ordinary one-way ANOVA with Tukey’s multiple comparisons test; ns, non-significant (*P* > 0.05), \*\*\*\**P* ≤ 0.0001. **(B)** Representative immunoblots showing the levels of IP6K1 and RAD51 knockdown in U-2 OS DR-GFP cells for data shown in (A); Data (mean ± SEM, *N* = 3) show the extent of IP6K1 and RAD51 knockdown quantified by normalising their respective band intensities to GAPDH and α-TUBULIN used as loading controls. (C) ChIP-qPCR analysis quantifying the relative enrichment of overexpressed SFB (vector control), or SFB-tagged RAD51 and IP6K1 near the I-SceI-generated DNA break at positions P1, P2, and P3 (see Supplementary Figure S6B). ChIP-qPCR data for each sample was normalised to its respective input levels. Results are presented as fold enrichment upon I-SceI-induced DNA damage relative to the respective uninduced sample. Data (mean ± SEM, *N* = 4), were analyzed using ordinary one-way ANOVA with Tukey’s multiple comparisons test; \**P* ≤ 0.05, \*\**P* ≤ 0.001, \*\*\*\**P* ≤ 0.0001. **(D, E, and F)** Representative immunoblots examining co-immunoprecipitation of endogenous RAD51 (D) (*N* = 2); overexpressed myc-BRCA2-CTD (E) (*N* = 2), or endogenous CDK1 (F) (*N* = 2) with endogenous IP6K1 in HEK293T lysates prepared from either untreated cells or MMC treated cells. IgG was used as a control. **(G, H, and I)** Representative immunoblots examining pull-down (PD) of endogenous RAD51 (G) (*N* =3), overexpressed myc-BRCA2-CTD (*N* = 3) (H), and endogenous CDK1 (*N*=3) (I) with either SFB-tagged active or inactive (K226A/S334A) mouse IP6K1 expressed in HEK293T cells, following treatment with MMC. The SFB-tagged proteins were pulled down with Streptavidin Sepharose beads (which bind the Streptavidin-binding peptide, SBP) and probed with an anti-FLAG antibody.

BRCA2, a large multi-domain protein, is a key regulator of RAD51 recruitment and removal during homology directed DNA repair [42, 46–48]. The C-terminal domain (CTD) of BRCA2, which can independently interact with RAD51 oligomers, is known to stabilize the RAD51 nucleoprotein filament and facilitate DNA strand invasion [49, 50, Kwon, 2023 #159, Kwon, 2023 #159, Kwon, 2023 #159, Kwon, 2023 #159]. CDK1-mediated phosphorylation of a conserved Ser residue in BRCA2-CTD lowers its binding with RAD51 to promote dissolution of RAD51 complexes [42, 46, 49]. Substitution of this Ser residue with either Ala to prevent phosphorylation, or with Glu to mimic constitutive phosphorylation, disrupt the binding of BRCA2-CTD with RAD51 and promote RAD51 foci disassembly, but do not impact HR efficiency [42, 46, 51]. We wondered whether IP6K1 binding to DNA breaks near RAD51 influences the interplay between RAD51, BRCA2-CTD, and CDK1 in the regulation of foci disassembly. We first examined the interaction of IP6K1 with these three proteins. IP6K1 immunoprecipitated with an antibody directed against its central region was able to co-precipitate endogenous RAD51, irrespective of DNA damage by MMC (Figure 5D and Supplementary Figure S6C). An antibody recognising the N-terminal region of IP6K1 [28] co-precipitated overexpressed BRCA2-CTD and endogenous CDK1 in untreated and MMC treated cells (Figure 5E, F and Supplementary Figure S6C). We then tested whether 5-InsP_7_ synthesis by IP6K1 influences its interaction with RAD51, CDK1 and BRCA2-CTD. SFB-tagged versions of both active and inactive IP6K1 were able to pull down RAD51 and BRCA2-CTD, indicating that the synthesis of 5-InsP_7_ does not influence the binding of IP6K1 to these proteins (Figure 5G, H and Supplementary Figure S6D). Interestingly, CDK1 interaction with inactive IP6K1 was significantly lower than its interaction with active IP6K1 (Figure 5I and Supplementary Figure S6D). This data, along with our earlier observation that CDK1-CCNB1 phosphorylates IP6K1 and upregulates its activity (Figure 3G - I), suggests that the interaction between IP6K1 and CDK1 occurs in a positive feedback loop, with CDK1 activating IP6K1 to synthesize 5-InsP_7_, which further promotes IP6K1-CDK1 binding.

To investigate whether the interaction between IP6K1 and these DNA repair proteins is direct or indirect, we prepared recombinant proteins expressed in *E. coli*. Immobilized GST-tagged RAD51, CDK1, or BRCA2-CTD were incubated with purified hexahistidine-tagged IP6K1. We observed direct binding of IP6K1 with both BRCA2-CTD and CDK1, but not with RAD51 (Supplementary Figure S6E). However, direct interaction of hexahistidine-tagged RAD51 was detected with both BRCA2-CTD and CDK1 (Supplementary Figure S6F). We also confirmed that RAD51 interacts with both BRCA2-CTD and CDK1 in cells treated with MMC (Supplementary Figure S6G and H). These protein-protein interaction data reveal that IP6K1 binds directly with BRCA2-CTD and CDK1, and indirectly, perhaps via these two proteins, with RAD51 (Supplementary Figure S6I).

### 5-InsP7 synthesis by IP6K1 disrupts the interaction between RAD51 and BRCA2-CTD

Next, we examined the effect of IP6K1 on the binding between RAD51 and BRCA2-CTD. When GST-tagged recombinant RAD51 purified from *E. coli* was allowed to interact with extracts from wild type or *IP6K1* KO HEK293T cells expressing BRCA2-CTD, we observed no difference in the extent of BRCA2-CTD and RAD51 binding (Figure 6A and B). Conversely, GST-tagged recombinant BRCA2-CTD expressed in *E. coli* showed a significantly higher degree of binding to endogenous RAD51 present in *IP6K1* KO U-2 OS cells compared with control cells (Figure 6C and D). The same effect was observed in HEK293T cells, where RAD51 in lysates from *IP6K1* KO cells showed enhanced binding with GST-BRCA2-CTD compared with RAD51 from wild type cells (Figure 6E and F). Adding back catalytically active IP6K1 in *IP6K1* KO cells lowered the interaction of RAD51 with GST-BRCA2-CTD, but inactive IP6K1 did not have the same effect (Figure 6E and F). These data reveal that 5-InsP_7_ synthesis by IP6K1 diminishes the interaction between BRCA2-CTD and RAD51. Additionally, 5-InsP_7_ affects this interaction when RAD51 is expressed in mammalian cells where it is capable of undergoing post-translational modifications, and BRCA2-CTD is expressed in *E. coli* where it is devoid of any post-translational modifications, but not vice versa. These findings indicate that the target of 5-InsP_7_ in the modulation of RAD51-BRCA2-CTD binding is RAD51 and not BRCA2-CTD.

**Figure 6.**
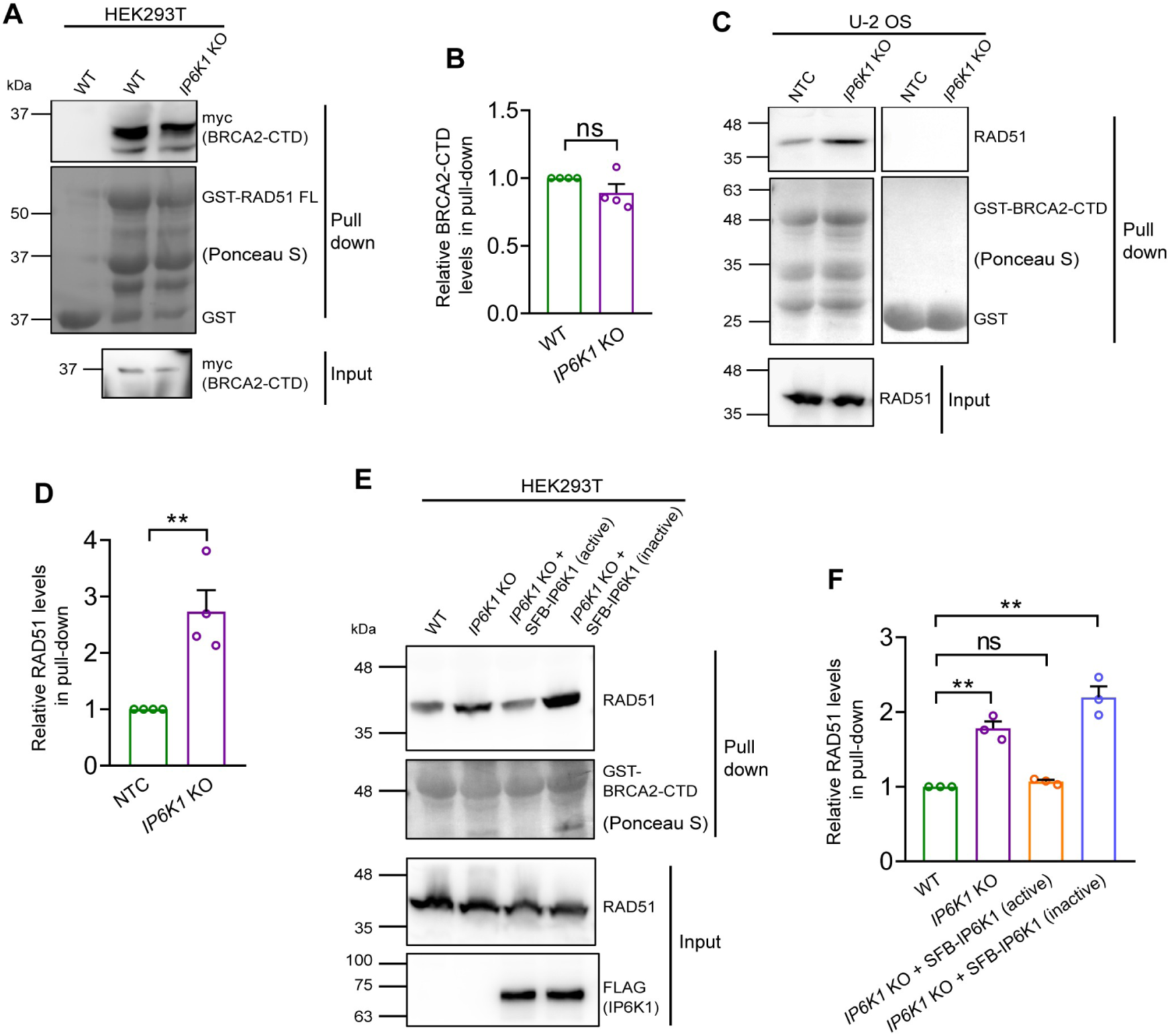
IP6K1 catalytic activity influences RAD51 binding with BRCA2-CTD. **(A)** Representative immunoblots examining interaction of myc-tagged BRCA2-CTD overexpressed in HEK293T WT and *IP6K1* knockout (KO) cells, with GST-tagged full length (FL) RAD51. **(B)** Quantification of (A), showing the relative extent of pull-down of BRCA2-CTD with GST-RAD51 in *IP6K1* KO compared with HEK293T WT cells, normalised to their respective input levels. Data (mean ± SEM, *N* = 4) were analyzed using a one-sample *t* test; ns, non-significant (*P* > 0.05). **(C)** Representative immunoblots examining interaction of endogenous RAD51 from MMC treated U-2 OS NTC and *IP6K1* knockout (KO) cells, with GST-tagged BRCA2-CTD. **(D)** Quantification of (C), showing the relative extent of pull-down of RAD51 with GST-BRCA2-CTD in *IP6K1* KO compared with NTC U-2 OS cells, normalised to their respective input levels. Data (mean ± SEM, *N* = 4) were analyzed using a one-sample *t* test; \*\**P* ≤ 0.01. **(E)** Representative immunoblots examining interaction of endogenous RAD51 from MMC treated HEK293T WT, *IP6K1* KO, and *IP6K1* KO cells overexpressing SFB-tagged active or inactive (K226A/S334A) mouse IP6K1, with GST-tagged BRCA2-CTD. **(F)** Quantification of (E), showing the relative extent of pull-down of RAD51 with GST-BRCA2-CTD in the indicated cell lines compared with HEK293T WT cells, normalised to their respective input levels. Data (mean ± SEM, *N* = 3), were analyzed using a one-sample *t* test; ns, non-significant (*P* > 0.05), \*\**P* ≤ 0.01. The GST-tagged proteins in (A, C, and E) were expressed in *E.coli*, and immobilised on glutathione Sepharose beads prior to use in the assay, and are visualised by staining with Ponceau S. GST, used as a control is shown in (A and C).

### 5-InsP7-mediated pyrophosphorylation at the N-terminus of RAD51 facilitates its dissociation from BRCA2-CTD

5-InsP_7_ can modulate protein-protein interactions by binding to a target protein or bringing about its pyrophosphorylation [1, 2, 9, 11]. To distinguish between these possibilities, one can employ 5-PCP-InsP_5_, a non-hydrolysable analog of 5-InsP_7_ that can bind to proteins at the same site as 5-InsP_7_ but cannot transfer its β phosphate [15, 52]. InsP_6_ has also been shown to bind to the same sites on many proteins as 5-InsP_7_ [52–54]. We therefore tested the impact of InsP_6_, 5-InsP_7_, and 5-PCP-InsP_5_ on the interaction between RAD51 and BRCA2-CTD. We observed no effect of pre-incubation of InsP_6_, 5-InsP_7_, or 5-PCP-InsP_5_ with GST-RAD51 on its interaction with BRCA2-CTD (Figure 7A and B). As InsP_6_ and 5-PCP-InsP_5_ can bind to proteins at the same site as 5-InsP_7_, this data suggests that the effect of 5-InsP_7_ on the RAD51-BRCA2-CTD interaction is not via binding, but may instead be via pyrophosphorylation.

**Figure 7.**
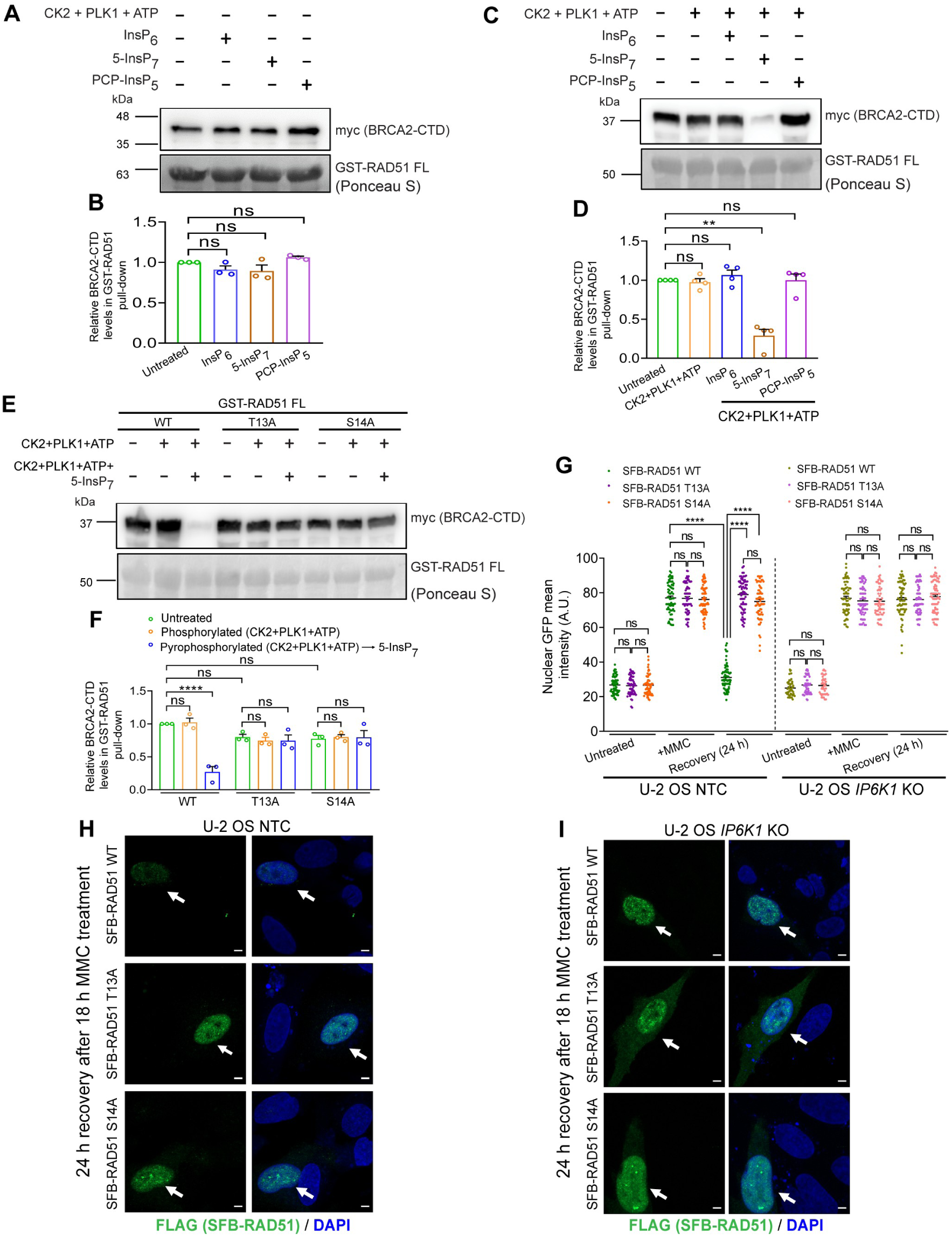
Effect of InsPs on RAD51 interaction with BRCA2-CTD. **(A-D)** Representative immunoblots examining the effect of pre-incubation with InsP_6_, 5-InsP_7_, and 5-PCP-InsP_5_ on the ability of unphosphorylated (A) or CK2 and PLK1 phosphorylated (C) GST-RAD51 to pull down myc-tagged BRCA2-CTD from MMC treated HEK293T cell lysates. Quantification of (A) and (C) show the extent of BRCA2-CTD bound to GST-RAD51 following treatment with the indicated InsP without (B) or with (D) pre-phosphorylation by CK2 and PLK1, normalised to untreated GST-RAD51. **(E)** WT or mutant (T13A or S14A) GST-RAD51 were left untreated, pre-phosphorylated with CK2 and PLK1, or incubated with 5-InsP_7_ subsequent to pre-phosphorylation. Representative immunoblots show the extent of myc-tagged BRCA2-CTD bound to GST-RAD51 in each case. **(F)** Quantification of (E) showing the extent of BRCA2-CTD bound to GST-RAD51 in each case, normalised to untreated WT GST-RAD51. Data (mean ± SEM; *N* = 3 in (B), *N* = 4 in (D), and *N* = 3 in (F)) were analyzed using one-way ANOVA with Tukey’s multiple comparisons test; ns, non-significant (*P* > 0.05), \*\**P* ≤ 0.01, \*\*\*\**P* ≤ 0.0001. GST-RAD51 in (A, C, and E) was expressed in *E.coli*, and immobilised on glutathione Sepharose beads prior to use in the assay, and is visualised by staining with Ponceau S. **(G-I)** Effect of IP6K1 on nuclear retention of RAD51 mutants upon MMC treatment and recovery. Representative immunofluorescence images of U-2 OS NTC and *IP6K1* KO cells overexpressing SFB-tagged RAD51 WT, T13A, or S14A, that were left untreated (Supplementary Figure S8A), treated with MMC (Supplementary Figure S8B), or allowed to recover from MMC-induced DNA damage for 24 h (H, I), stained with anti-FLAG antibody (green) to detect the SFB-tag. The transfected cells are indicated with white arrows, nuclei were stained with DAPI (blue), and scale bars are 5 μm. Quantification (G) of SFB-RAD51 nuclear intensity in (H, I), S8A and S8B. Data distribution is represented by scatter dot plots (*n* = 64, 60, and 62 for SFB-RAD51 WT, T13A, and S14A expressed in untreated U-2 OS NTC cells; *n* = 69, 64, and 70 for SFB-RAD51 WT, T13A, and S14A in MMC treated U-2 OS NTC cells; *n* = 77, 70, and 61 for SFB-RAD51 WT, T13A, and S14A expressed in post-recovery U-2 OS NTC cells; *n* = 47, 51, and 42 for SFB-RAD51 WT, T13A, and S14A expressed in untreated U-2 OS *IP6K1* KO cells; *n* = 63, 65, and 59 for SFB-RAD51 WT, T13A, and S14A in MMC treated U-2 OS *IP6K1* KO cells; and *n* = 70, 70, and 54 for SFB-RAD51 WT, T13A, and S14A expressed in post-recovery U-2 OS *IP6K1* KO cells). Data are compiled from four independent experiments, and were analyzed using Kruskal-Wallis test with Dunn’s multiple comparison post hoc test; ns, non-significant (*P* > 0.05), \*\*\*\**P* ≤ 0.0001.

To explore the possibility of RAD51 pyrophosphorylation by 5-InsP_7_, we first needed to identify the protein kinase(s) that catalyse the priming phosphorylation pre-requisite for pyrophosphorylation. RAD51 has previously been reported to undergo phosphorylation by CK2 and PLK1 [41]. Like CK2, which is known to pre-phosphorylate proteins to prime them for pyrophosphorylation, PLK1 is also an acidophilic protein kinase, and recognises substrate motifs containing Asp/Glu at the −2 position with respect to the target Ser/Thr residue [55]. As CDK1-CCNB1 phosphorylates both IP6K1 (Figure 3G) and BRCA2-CTD [42], we also tested whether it is capable of phosphorylating RAD51. In the presence of radiolabelled ATP, RAD51 underwent robust phosphorylation by CK2 and PLK1, but was not phosphorylated by CDK1-CCNB1 (Supplementary Figure S7A). To map the CK2 and PLK1 phosphosites on RAD51, we generated GST-fusion fragments encompassing different RAD51 domains (Supplementary Figure S7B). RAD51 consists of an N-terminal DNA binding domain, a linker region, and a globular C-terminal ATPase domain [56]. The N-terminal domain of RAD51 (residues 1-98), was phosphorylated by both CK2 and PLK1, but the C-terminal region (residues 99-339) was not phosphorylated by either protein kinase (Supplementary Figure S7C and D). Structural studies on RAD51 in its monomeric or DNA-bound forms have revealed that ∼20 residues at the N-terminus of RAD51 are unstructured [57, 58]. RAD51 constructs lacking this disordered N-terminus did not undergo phosphorylation by either CK2 or PLK1 (Supplementary Figure S7C and D), narrowing down the region of phosphorylation to this stretch of 20 residues. The only potential sites for phosphorylation in this sequence are Thr13, Ser14 and Ser19. An earlier study had identified Ser14 as the site for phosphorylation on RAD51 by PLK1, which enables subsequent phosphorylation on Thr13 by CK2 [41]. We conducted site-directed mutagenesis on full length GST-tagged RAD51 to replace Thr13, Ser14, or Ser19 with Ala, and examined phosphorylation of these mutants by CK2 and PLK1. Substitution of Thr13 eliminated CK2-mediated phosphorylation on RAD51, whereas the Ser14Ala mutant showed reduced phosphorylation (Supplementary Figure S7E). Conversely, mutation of Ser14 to Ala prevented RAD51 phosphorylation by PLK1, and the Thr13Ala mutant showed reduced phosphorylation (Supplementary Figure S7F). Replacing Ser19 with Ala had no impact on phosphorylation by either CK2 or PLK1 (Supplementary Figure S7E and F). In the case of the RAD51 (1-98), the Thr13Ala mutant was phosphorylated by PLK1 but not by CK2, and the Ser14Ala mutant was phosphorylated by CK2 and not by PLK1 (Supplementary Figure S7G and H). These data confirm that CK2 and PLK1 phosphorylate Thr13 and Ser14 respectively, and that Ser 19 is not a target for either of these protein kinases.

5-InsP_7_-mediated protein pyrophosphorylation is known to preferentially occur on Ser residues that lie close to Asp/Glu residues in regions of intrinsic disorder [8, 17]. As the phosphorylated residues (Ser13 and Thr14) at the N-terminus of RAD51 are located within a disordered sequence [57, 58], we examined whether the RAD51 N-terminal domain undergoes pyrophosphorylation. We pre-phosphorylated GST-RAD51 (1-98) with CK2 or PLK1 individually or together in the presence of unlabelled ATP, and subsequently incubated the protein with radiolabelled 5-InsP_7_ (Supplementary Figure S7I). RAD51(1-98) accepted radiolabelled β-phosphate from 5-InsP_7_ subsequent to phosphorylation by CK2, PLK1, or both, indicating that Thr13 and Ser14 are capable of undergoing pyrophosphorylation (Supplementary Figure S7J). This is the first time that PLK1 catalysed phosphorylation has been shown to prime a Ser residue for pyrophosphorylation. We then examined the effect of substituting Thr13 or Ser14 with Ala on 5-InsP_7_-mediated pyrophosphorylation of RAD51. The wild type and point mutant versions of RAD51(1-98) were pre-phosphorylated with a mix of CK2 and PLK1, and then incubated with radiolabelled 5-InsP_7_. Interestingly, mutation of either Thr13 or Ser14 to Ala nearly eliminated pyrophosphorylation on the RAD51 N-terminal domain (Supplementary Figure S7K), indicating that both residues are essential for pyrophosphorylation to occur on either residue.

Having established that RAD51 can undergo 5-InsP_7_-mediated pyrophosphorylation, we monitored the impact of this modification on the interaction of RAD51 with BRCA2-CTD. Phosphorylation of GST-tagged full length RAD51 by CK2 and PLK1 had no effect on its binding with BRCA2-CTD compared with the unphosphorylated control (lanes 1 and 2 in Figure 7C and D). Subsequent incubation of phosphorylated GST-RAD51 with either InsP_6_ or 5-PCP-InsP_5_ also did not alter the binding of RAD51 with BRCA-CTD (lanes 3 and 5 in Figure 7C and D). In contrast, incubation with 5-InsP_7_, which is proficient in pyrophosphorylation, significantly attenuated the binding of RAD51 with BRCA2-CTD (lane 4 in Figure 7C and D). 5-InsP_7_ did not have the same effect on Thr13Ala or Ser14Ala RAD51 mutants. When these GST-RAD51 mutants were pre-incubated with CK2 and PLK1 in the presence of ATP, further addition of 5-InsP_7_ did not alter their binding with BRCA2-CTD (Figure 7E and F). Together, these data show that mere phosphorylation of RAD51 on Thr13 and Ser14 does not impact its binding with BRCA2-CTD, but that pyrophosphorylation on these residues is required to disrupt BRCA2-CTD binding to RAD51.

Finally, we assessed the nuclear accumulation of wild type and Thr13Ala or Ser14Ala mutant forms of RAD51 during MMC-induced DNA damage and recovery. Upon treatment with MMC, full length SFB-tagged RAD51 accumulated in the nucleus in both U-2 OS NTC and *IP6K1* KO cells (Figure 7G, Supplementary Figure S8A and B). In U-2 OS NTC cells, as expected, we observed reduced nuclear intensity of SFB-RAD51 when cells were allowed to recover for 24 h after removal of MMC (Figure 7G and H). However, in *IP6K1* KO U-2 OS cells, SFB-RAD51 remained in the nucleus 24 h after MMC removal (Figure 7G and I), mirroring the behaviour of endogenous RAD51 in these cells (Supplementary Figure S2 A and B). We confirmed that there was no change in the levels of SFB-RAD51 in both cell lines during MMC treatment and recovery (Supplementary Figure S8C and D). Thr13Ala and Ser14Ala SFB-RAD51 mutants expressed in U-2 OS NTC cells showed nuclear recruitment upon MMC treatment similar to wild type SFB-RAD51 (Figure 7G, Supplementary Figure S8A and B). This data is in contrast to an earlier report which showed that loss of phosphorylation on Ser14 led to a reduction in the formation of RAD51 nuclear foci upon irradiation-induced DNA damage [41]. Unlike wild type SFB-RAD51, the mutant forms of RAD51 expressed in U-2 OS NTC cells retained their nuclear localisation even after removal of MMC (Figure 7G and H). As in the case of wild type SFB-RAD51, Thr13Ala and Ser14Ala mutants showed nuclear retention in *IP6K1* KO U-2 OS cells following MMC removal (Figure 7G and I). Taken together, these data provide convincing evidence that 5-InsP_7_-mediated pyrophosphorylation of RAD51 on Thr13 and Ser14 promotes its dissociation from BRCA2-CTD and leads to dissolution of RAD51 nuclear foci post-repair.

## Discussion

Our study underlines the pivotal role played by the inositol pyrophosphate 5-InsP_7_ in enabling the removal of RAD51 from DNA damage sites. Building on our earlier demonstration that IP6K1 knockout mouse embryonic fibroblasts display persistent RAD51 foci during recovery from genotoxic stressors [22], we now show that 5-InsP_7_ synthesised by IP6K1 pyrophosphorylates the N-terminus of RAD51, promoting the dissociation of RAD51 from the C-terminus of BRCA2 to facilitate the dissolution of DNA damage foci (Figure 8). Cells depleted for 5-InsP_7_ show a reduction in their ability to disassemble RAD51 foci during recovery from treatment with MMC. 5-InsP_7_ synthesis by IP6K1 increases upon MMC-induced DNA damage, and this is attributed to phosphorylation of IP6K1 in its N-lobe by CK2 and CDK1-CCNB1. IP6K1 localises near the site of DNA damage, and engages in an array of interactions with proteins known to be present at the damage site, including RAD51, CDK1, and the C-terminal domain of BRCA2. The presence of IP6K1 near the DNA break is likely to increase the local concentration of 5-InsP_7_ and promote pyrophosphorylation of RAD51 at Thr13 and Ser14, subsequent to its phosphorylation by CK2 and PLK1. This pyrophosphorylation on RAD51 attenuates its interaction with BRCA2-CTD, promoting the dissolution of RAD51 nuclear foci during recovery from DNA damage.

**Figure 8.**
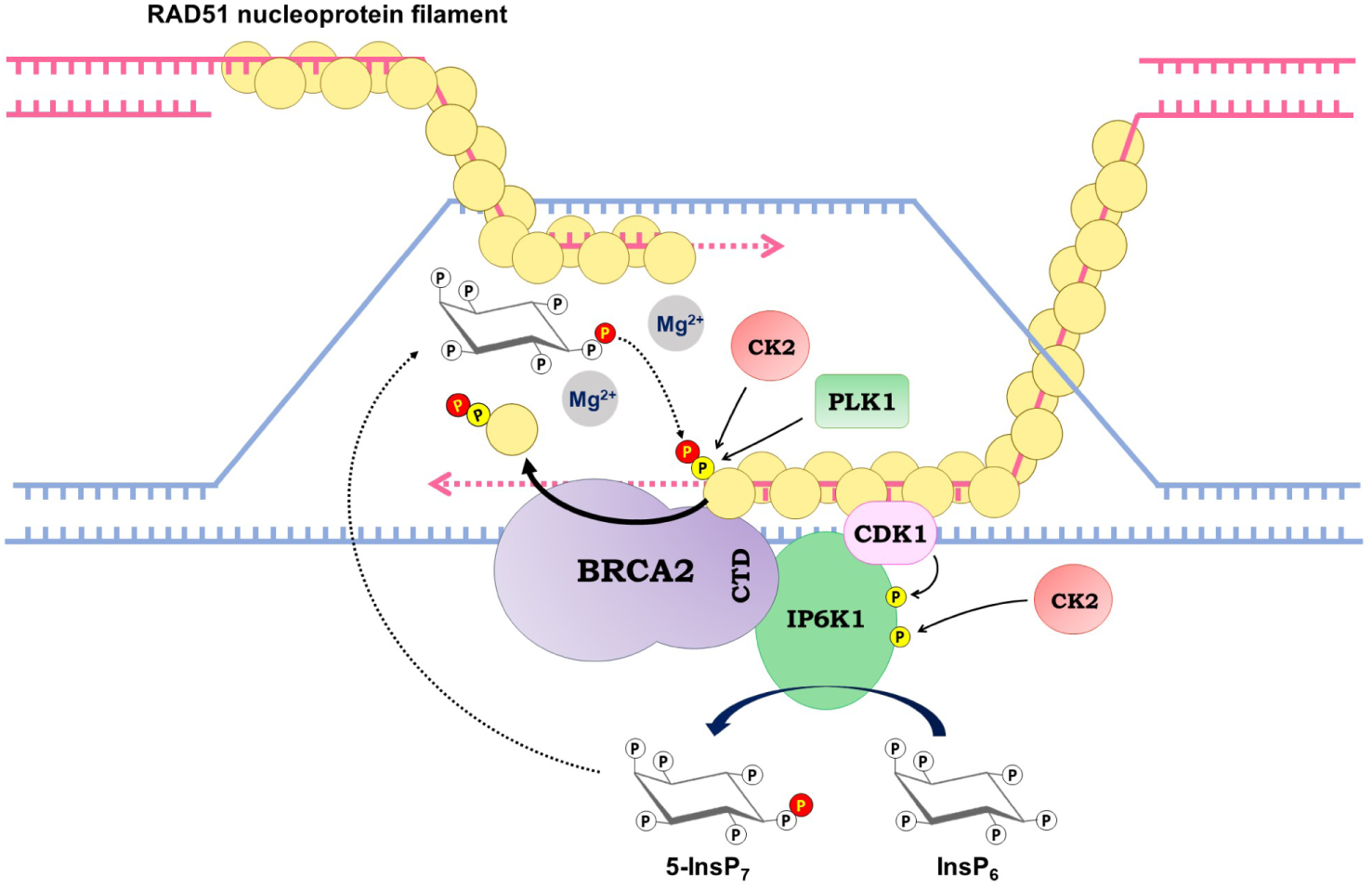
5-InsP_7_-mediated pyrophosphorylation promotes RAD51 removal from DNA damage foci. During HR-mediated repair, RAD51 loading onto ssDNA to form a nucleoprotein filament is aided by BRCA2. RAD51 is phosphorylated at its N-terminus by the acidophilic Ser/Thr kinases CK2 and PLK1. IP6K1, recruited near the double strand break, interacts with DNA repair proteins including CDK1 and the C-terminus (CTD) of BRCA2. Phosphorylation by CDK1 and CK2 promotes IP6K1 activity to increase the synthesis of 5-InsP_7_, which in the presence of Mg^2+^, pyrophosphorylates the N-terminus of RAD51. This modification on RAD51 reduces its interaction with BRCA2-CTD, leading to RAD51 eviction from DNA damage sites.

Pyrophosphorylation has been shown to preferentially occur on Ser residues located within regions of intrinsic disorder [8, 17]. In RAD51, one of the pyrophosphorylated residues is Thr, which we have shown in the case of the oncoprotein c-MYC, is a poorer acceptor of the β-phosphate from 5-InsP_7_ compared with Ser in the same sequence context [16]. Intrinsically disordered protein sequences that are sites for pyrophosphorylation are also hubs for protein-protein interaction. Earlier studies have shown that 5-InsP_7_-dependent pyrophosphorylation of a target protein can have a positive or negative impact on its interaction with other proteins [11]. Upon pyrophosphorylation by 5-InsP_7_, DIC2C (intermediate chain of the dynein motor complex) and the oncoprotein c-MYC exhibit augmented binding to the dynactin subunit p150*^Glued^* and the E3 ubiquitin ligase FBW7 respectively [15, 16]. On the other hand, pyrophosphorylation diminishes binding of the yeast glycolytic transcription factor GCR1, and the AP3 adaptor protein complex β subunit AP3B1, with their respective binding partners GCR2 and the kinesin motor protein Kif3A [13, 59]. Our demonstration that RAD51 pyrophosphorylation promotes its dissociation from BRCA2-CTD provides another example of disruption of protein-protein interaction by pyrophosphorylation, where phosphorylation alone has no impact. Phosphorylation by protein kinases is a key regulatory mechanism in DNA repair [60]. Our work reveals that CK2 and CDK1-CCNB1-mediated phosphorylation in the N-lobe of IP6K1 augments its enzyme activity upon DNA damage (Figure 3H and I). CK2 phosphorylation on IP6K1 occurs within a 12 residue stretch (Ser118-Ser129) which contains 5 Ser residues (Figure 4C and S5D). Ser118 and Ser121 have previously been shown to undergo phosphorylation by protein kinases PKA, PKC and PKD, leading to upregulation of IP6K1 activity [19, 61]. As reported earlier, we have confirmed phosphorylation of RAD51 on Thr13 and Ser14 by CK2 and PLK1 respectively [41]. Ser14 phosphorylation by PLK1 is known to licence RAD51 for subsequent phosphorylation by CK2 on Thr13, promoting RAD51 binding with NBS1 (Nijmegen breakage syndrome) to facilitate its recruitment to DNA damage sites [41]. Interestingly, our work reveals a contrasting molecular impact of pyrophosphorylation on the same residues – whereas phosphorylation on Thr13 and Ser14 promotes RAD51 recruitment to DNA breaks, pyrophosphorylation on these residues facilitates RAD51 removal from DNA damage sites.

The interaction of RAD51 with BRCA2 plays a key role in RAD51 nucleoprotein filament dynamics during DNA damage repair. RAD51 loading onto ssDNA to form a nucleoprotein filament is supported by its interaction with BRC repeats in BRCA2 [47, 48]. The CTD of BRCA2, which is unrelated in sequence to BRC repeats, also plays a crucial role in loading RAD51 oligomers onto ssDNA, and deletion of this BRCA2 domain results in impaired HR efficiency and reduced survival in mice [50, Kwon, 2023 #159, Kwon, 2023 #159, Kwon, 2023 #159, Kwon, 2023 #159, 62, 63]. The loss of interaction between BRCA2-CTD and RAD51 is known to promote the dissociation of RAD51 from DNA at repair sites, leading to the dissolution of RAD51 nuclear foci [42, 49, 50]. Phosphorylation of BRCA2-CTD by CDK1 has been shown to disrupt the association of BRCA2 with RAD51, supporting the disassembly of damage-induced RAD51 foci [42, 46]. Point mutations in BRCA2-CTD that reduce its interaction with RAD51 are dispensable for HR, but lead to faster dissolution of RAD51 foci during recovery from DNA damage [42, 46, 51]. These reports are in congruence with our findings that RAD51 pyrophosphorylation, which disrupts its binding with BRCA2-CTD, supports RAD51 foci dissolution. Our data showing that depletion of IP6K1 increases RAD51 binding with BRCA2-CTD, but does not impact HR, is also in line with these previous studies. In summary, our demonstration that RAD51 pyrophosphorylation promotes dissociation of BRCA2-CTD from RAD51 adds a new dimension to the regulatory landscape governing RAD51 removal from damage sites post DNA repair.

## Methods

Information on reagents and routine methods is provided in the Supplementary Methods.

### Analysis of cellular inositol pyrophosphates

Pulldown of inositol phosphates on titanium dioxide beads and PAGE analysis was performed as described previously [64]. Briefly, cells in 15 cm dishes (4 x 15 cm dishes for U-2 OS and 2 x 15 cm dishes for HEK293T) were treated with 10 mM NaF for 1.5 h, followed by trypsinization. Inositol pyrophosphates were extracted using 1 M perchloric acid (HClO_4_). The extracts were incubated with 5-7 mg titanium dioxide beads (Titansphere TiO 5 μm; GL Sciences) per sample. Inositol phosphates were eluted by the addition of ∼2.8% ammonia water (NH_4_OH; 2 x 200 µl). The eluates were reduced in volume by centrifugal evaporation until neutral pH was attained (down to ∼40 µl), were resolved using 35.8% polyacrylamide gels, and visualized by Toluidine blue staining. The gels were scanned on a desktop scanner. For [^3^H]inositol labelling and SAX-HPLC analysis, HEK293T or U-2 OS cells were seeded in 60 mm dishes and incubated in inositol-free DMEM (MP Biomedicals) containing 10% dialysed fetal bovine serum. Cells were labelled with 30 µCi or 40 µCi *myo*-2-[^3^H] inositol for HEK293T or U-2 OS cell lines respectively, and incubated for 2.5 days. The media was removed and fresh media containing the same amounts of *myo*-2-[^3^H] inositol was added for another 2.5 days. The inositol phosphates were resolved by HPLC and quantified as described earlier [17, 30, 65]. Soluble inositol phosphates were extracted from cells by incubation with 350 µL 0.6 M HClO_4_, 2 mM EDTA, and 0.2 mg/mL phytic acid for 20 min at 4°C, followed by centrifugation at 21,000 x *g* for 10 min. The supernatant containing soluble inositol phosphates was collected, and inositol containing lipids in the pellet were solubilized in 0.1 N NaOH, 0.1% Triton X-100 at room temperature (RT), and counted in a liquid scintillation counter (Tri-Carb 2910 TR, PerkinElmer). The soluble inositol phosphate extract was mixed with ∼120 µL neutralization solution (1 M K_2_CO_3_, 5 mM EDTA). Tubes were left open on ice for 1 h, followed by centrifugation at 21,000 x *g* for 10 min at 4 °C. The extracted inositol phosphates were resolved by HPLC (Waters 515 or 5125 HPLC pumps) on a PartiSphere SAX column (4.6 mm × 125 mm, HiChrome) using a gradient of buffer A (1 mM EDTA) and buffer B (1 mM EDTA and 1.3 M (NH_4_)_2_HPO_4_ (pH 3.8)) as follows: 0–5 min, 0% B; 5–10 min, 0–20% B; 10–70 min, 20–100% B; 70–80 min, 100% B. 1 mL fractions containing soluble inositol phosphates were mixed with 3 mL of scintillation cocktail (Ultima-Flo AP) and counted.

### IP6K1 enzyme activity assay

To determine the activity of overexpressed GFP-tagged catalytically active and inactive IP6K1, and immunoprecipitated endogenous IP6K1, the enzymes immobilised on Protein A Sepharose beads were incubated with 200 µM InsP_6_, 50 µM Mg^2+^-ATP, reaction buffer (20 mM MES, pH 6.4, 50 mM NaCl, 6 mM MgSO_4_, 1 mM DTT) in the presence of 3 µCi [γ-^32^P]ATP at 37°C for 1 h. The reactions were terminated by the addition of 3 mM EDTA, and 75 µL 0.6 M perchloric acid (HClO_4_) was added to each tube. The reactions were neutralised with ∼33 µL 1 M K_2_CO_3_ and 5 mM EDTA, and centrifuged at 21,000 x *g* for 15 min at RT. The clarified extracts were resolved by SAX-HPLC as described above, and 1 mL fractions corresponding to the 5-InsP_7_ peak were counted.

### Assessment of HR-mediated repair

The U-2 OS DR-GFP (mCherry-I-SceI-GR) cell line (a gift from Dr. Evi Soutoglou, IGBMC, Strasbourg, France) was maintained in DMEM without phenol red (Thermo Fisher Scientific) supplemented with 10% charcoal stripped fetal bovine serum (Thermo Fisher Scientific), 1 mM L-glutamine, 100 U/mL penicillin, 100 µg/mL streptomycin, and 1 mM sodium pyruvate (Thermo Fisher Scientific), supplemented with 1 µg/mL puromycin and 400 µg/mL G418. HR-mediated repair was assessed as described earlier [66, 67]. Transient knockdown was achieved by transducing cells with lentiviral particles encoding shNT, sh*IP6K1*, or sh*RAD51*. For the generation of lentiviral particles, HEK293T cells were co-transfected with lentiviral vectors (pLKO.1) encoding non-targeting control shRNA (shNT) or shRNA directed against human *IP6K1* or *RAD51*, and packaging plasmids VSV-G and psPAX2, using polyethylenimine (PEI). Transfected HEK293T cells were incubated in phenol red-free DMEM supplemented with 10% charcoal stripped fetal bovine serum at 37°C for 48 h. Lentiviral particles were harvested by filtering the culture supernatant through a 0.45 μm filter. U-2 OS DR GFP cells were treated with 8 μg/mL polybrene for 2 h prior to lentiviral transduction. 72 h post-transduction, cells were provided fresh media containing either 1% DMSO (vehicle control) or 0.1 µM triamcinolone acetonide (TA) to induce I-SceI-mediated DNA damage. After 72 h, cells were harvested by trypsinisation and analyzed for GFP expression by flow cytometry (BD Accuri C6). A parallel set was used to determine the extent of knockdown by western blotting.

### ChIP-on-break assay

ChIP-qPCR assays were performed in U-2 OS DR-GFP (mCherry-I-SceI-GR) cells to detect the occupancy of RAD51 or IP6K1 near the I-SceI-induced DNA break. Cells overexpressing SFB-tagged IP6K1 or RAD51 were incubated with 0.1 μM triamcinolone acetonide (TA) for 4 h to induce DNA double strand breaks. The cells were fixed with 1% formaldehyde in culture medium for 15 min at RT, followed by quenching in 0.125 M glycine for 10 min. Cells were lysed in buffer containing 50 mM HEPES-KOH pH 7.5, 140 mM NaCl, 1 mM EDTA, 1% Triton X-100, 0.1% SDS, 0.1% sodium deoxycholate and protease inhibitor cocktail, and sonicated to fragment chromatin to the desired extent. Cell lysates were diluted in immunoprecipitation dilution buffer (50 mM Tris-HCl pH 8.0, 150 mM NaCl, 2 mM EDTA, 1% Nonidet P-40, 0.5% sodium deoxycholate, 0.1% SDS, and protease inhibitor cocktail), and immunoprecipitation was carried out with anti-FLAG (M2) antibody overnight at 4°C, followed by Protein G Sepharose beads for 2 h. Beads were washed sequentially, twice each with low salt immune complex buffer (20 mM Tris-HCl pH 8.0, 150 mM NaCl, 2 mM EDTA, 0.1% SDS, 1% Triton X-100), high salt immune complex buffer (20 mM Tris-HCl pH 8.0, 500 mM NaCl, 2 mM EDTA, 0.1% SDS, 1% Triton X-100, and), LiCl buffer (10 mM Tris-HCl pH 8.0, 0.25M LiCl, 1 mM EDTA, 1% Nonidet P-40, 1% sodium deoxycholate,), and TE buffer (10 mM Tris-HCl pH 8.0, 1 mM EDTA). Immunoprecipitated chromatin was eluted with elution buffer (50 mM Tris-HCl pH 8.0, 10 mM EDTA, 0.5% SDS) and incubated for 16 h at 65°C to reverse the crosslinks. DNA was purified using standard phenol, chloroform, and isoamyl alcohol based extraction and ethanol precipitation. The relative occupancy of the immunoprecipitated proteins at each DNA locus was estimated by RT-qPCR. The primers for ChIP-qPCR analysis are listed in Supplementary Table S5.

### Protein phosphorylation and pyrophosphorylation

The synthesis of radiolabelled 5[β-^32^P]InsP_7_, protein phosphorylation and pyrophosphorylation was conducted as described previously [16, 17, 68, 69]. For phosphorylation assays, 2-4 µg of GST-tagged proteins immobilised on glutathione Sepharose beads were incubated with either CK2, CDK1-CCNB1, or PLK1 at 30°C in the presence of protein kinase buffer (50 mM Tris-HCl, pH 7.5, 15 mM MgCl_2_, 0.1 mM EDTA, 1 mM EGTA, 2 mM DTT) with 3 µCi [γ-^32^P]ATP, and 400 µM Mg^2+^-ATP. Beads were washed twice with ice-cold PBS prior to NuPAGE analysis. For pyrophosphorylation reactions, GST-tagged proteins immobilised on glutathione Sepharose beads were pre-phosphorylated by either CK2 or PLK1 or both protein kinases as above without the addition of [γ-^32^P]ATP. Beads were washed with ice-cold PBS, resuspended in pyrophosphorylation buffer (25 mM HEPES-KOH pH 7.4, 50 mM NaCl, 6 mM MgCl_2_, 1 mM DTT) in the presence of 7-10 µCi 5[β-^32^P]InsP_7_, and incubated at 37°C for 45 min. Following either phosphorylation or pyrophosphorylation, proteins were eluted by boiling the beads in 1X LDS sample buffer at 95°C for 5 min, resolved by 4-12% NuPAGE Bis-Tris gel, and transferred to a PVDF membrane. Blots were dried and exposed to a phosphorimager screen for 2-3 days for phosphorylation assays, and ∼30-40 days for pyrophosphorylation assays, and imaged using a phosphorimager (Typhoon FLA-9500). To improve visualization, phosphorimager scans were subjected to uniform ‘Levels’ adjustment in Adobe Photoshop. Ponceau S staining was used to detect the amount of protein loaded.

### Statistical analysis

GraphPad Prism 8 was used to perform statistical analyses and prepare graphs. The number of cells (n) used to perform statistical tests for each experiment, and the number of biologically independent replicates (N) for each experiment are indicated in the figure legends. Statistical significance was assessed by using appropriate parametric or non-parametric analyses as indicated in the respective figure legends. *P* ≤ 0.05 was considered statistically significant. For analysis of data presented as relative fold-change, individual groups were compared to a control that was normalized to 1. The values were log_2_-transformed to ensure normality and account for skewness of the data, and then subjected to a one-sample *t* test against a theoretical mean of 0; differences between multiple groups were assessed by one-way ANOVA followed by Tukey’s multiple comparisons test.

## Supplementary Information

Supplementary Methods and Supplementary Figures S1-S8 are provided in a single PDF file. Supplementary Tables S1-S7 are provided as Excel files.

## Data Availability Statement

The mass spectrometry data from this study are available on **MassIVE**, a Proteome Xchange Consortium member, and can be accessed through the dataset identifier **MSV000096311** at https://massive.ucsd.edu.

## Author Contributions

**Shubhra Ganguli**: Conceptualization, Methodology, Investigation, Validation, Formal analysis, Visualization, Writing – original draft, Writing – review and editing. **Rashna Bhandari**: Conceptualization, Writing – review and editing, Supervision, Project administration, Funding acquisition. All authors have read and agreed to the published version of the manuscript.

## Acknowledgements

We thank Dorothea Fiedler, Mingxuan Wu, and Leonie Kurz for generously sharing 5-InsP_7_ and 5-PCP-InsP_5_, and for helpful discussions; Sagar Sengupta for sharing pET28b RAD51 plasmid, and for helpful discussions; Ashok Venkitaraman for sharing pEFMN BRCA2-CTD and pGEX-4T3 BRCA2-CTD plasmids, and for valuable feedback; Evi Soutoglou for sharing the U-2 OS DR-GFP (mCherry-I-SceI-GR) cell line; P Chandra Shekar for sharing pU6-2A-GFP-2A-Puro plasmid; Ganesh Nagaraju for helpful discussions; Akruti Shah and Vineesha Oddi for generation of IP6K1 knockout U-2 OS and non-targeted control cell lines; Jayashree Suresh Ladke for generation of IP6K1 knockout HEK293T cell lines; the staff at the Sophisticated Equipment Facility for technical assistance; and members of the Laboratory of Cell Signalling for valuable feedback.

