## Supplementary Information for "The inositol pyrophosphate 5-InsP7 promotes DNA repair by disrupting RAD51 binding to the C-terminus of BRCA2"

**Running title**: 5-InsP$_7$ disrupts RAD51 – BRCA2 interaction

**ORCID iDs**

Shubhra Ganguli - 0000-0001-5519-592
Rashna Bhandari - 0000-0003-3101-0204

**This PDF file includes:**

Supplementary Materials and Methods

Legends for Supplementary Tables S1-S7

Supplementary Figures S1-S8

Supplementary References

**Supplementary Materials and Methods**

**Materials:** All chemicals were procured from Merck, unless specified otherwise. Restriction enzymes were purchased from New England Biolabs. Dulbecco's Modified Eagle's Medium (DMEM), Fetal Bovine Serum (FBS) and other cell culture reagents were from Thermo Fisher Scientific. NuPAGE 4-12% Bis-Tris gels, 20X MES running buffer, and 4X LDS sample buffer were purchased from Thermo Fisher Scientific. [$\gamma$-$^{32}$P]ATP (LCP-101) was procured from JONAKI/BRIT. *myo*-2-[$^{3}$H] inositol (15-20Ci/mmol) (ART 0116B) was procured from American Radiolabeled Chemicals. Ultima-Flo AP (6013599) was purchased from Perkin-Elmer. PVDF membrane for protein transfer, glutathione-Sepharose beads (17-0756-01), Protein A Sepharose beads (17-0780-01), Protein G Sepharose beads (17-0618-01), Streptavidin-Sepharose beads (17-5113-01), and ECL prime chemiluminiscence substrate were purchased from Cytiva Life Sciences. DNA damage agents used in the study, along with their concentration and duration of treatment, are listed in Supplementary Table S1. Protein kinases used for phosphorylation assays, along with the amount used in the study, are listed in Supplementary Table S2. Primary and secondary antibodies used in this study are listed in Supplementary Table S3. Plasmids used in this study are listed in Supplementary Table S4. Primers used for cloning, mutagenesis, and ChIP-on-break assay are listed in Supplementary Table S5, and shRNA and sgRNA sequences are listed in Supplementary Table S6.

**Cell culture, transduction, and transfection:** U-2 OS and HEK293T cell lines were grown in a humidified incubator with 5% $CO_2$ at 37°C in Dulbecco's modified Eagle's medium (DMEM) supplemented with 10% fetal bovine serum, 1 mM L-glutamine, 100 U/mL penicillin, and 100 µg/mL streptomycin. For transfection, polyethylenimine (PEI) (Polysciences, 23966) was used at a ratio of 1:3 (DNA:PEI). All plasmids used for transfection were purified using the Plasmid Midi kit (Qiagen). Cells were harvested 36-48 h post transfection for further analyses. All cell lines used tested free from mycoplasma. U-2 OS NT and sh*IP6K1* cell lines have been described earlier [1]. *IP6K1*$^{-/-}$ U-2 OS and HEK293T cell lines were generated using the CRISPR-Cas9-mediated knockout strategy. For U-2 OS cell lines, the sgRNA (single guide RNA) sequences (Supplementary Table S6) were designed by TransOMIC Technologies and provided in the pCLIP-ALL-EFS-Blasticidin destination vector. Lentiviral particles were packaged in HEK293T cells by co-transfecting plasmids encoding sgRNA with VSV-G and psPAX2. Post transfection, culture supernatants were collected and filtered to harvest viral particles, and U-2 OS cells were transduced following treatment with polybrene (8µg/mL). The cells were selected with 5 µg/mL blasticidin for 15 days. Single cells were derived by serial dilution in conditioned media, which were screened for frameshift mutations by genotyping using a 3500xL Genetic Analyzer (Applied Biosystems). The selected clones were expanded and screened for *IP6K1* knockout by western blotting analysis. The generation of *IP6K1*$^{-/-}$ HEK293T cell lines has been described previously [2].

**Immunofluorescence analysis:** Cells grown on glass coverslips were treated with DNA damage agents. Following incubation, the media was replaced and cells were allowed to recover for the indicated time. For γH2AX staining, coverslips were incubated in hypotonic lysis buffer (10 mM Tris-HCl pH 7.4, 2.5 mM $MgCl_2$, 0.5% Nonidet P-40, 1mM PMSF) for 8 min at 4°C, and were fixed with ice-cold 100% ethanol for 4 min. For IP6K1 and RAD51 staining, cells were fixed with freshly prepared 4% formaldehyde for 10 min, and permeabilised in 0.2% Triton X-100 for 10 min at room temperature (RT). Non-specific interactions were blocked by incubating the cells for 45-60 min with 3% BSA in 1X PBS. Coverslips were incubated with either γH2AX, IP6K1, or RAD51 antibody diluted in the blocking solution, overnight at 4°C in a moist chamber. Post incubation with primary antibodies, the coverslips were washed with PBS-T (PBS with 0.5% Tween 20), incubated with fluorophore conjugated secondary antibodies (Supplementary Table S3) for 1 h at RT. The coverslips were mounted on glass slides using an antifade mounting medium with DAPI (H-1200, Vecta Labs), air dried and sealed. Images were acquired using an LSM 700 or LSM 900 (Zen acquisition software) confocal microscope (Zeiss) equipped with 405, 488, and 555 nm lasers and fitted with a 63x, 1.4 N.A. objective. The exposure settings and other imaging parameters were identical for images of different cell types / treatment conditions in a single experiment. All images are in z-stacks and are shown as maximum intensity projections (MIP) using ZEN software. The number of γH2AX or RAD51 foci per cell was quantified using Fiji software [3]. Briefly, z-stacks (step size either 0.5 µm or 1 µm) were digitally collapsed, and then subtracted for the background (value 25). Foci were counted in each nucleus using the 'analyse particles' feature of Fiji. Pearson correlation coefficient analysis was performed using the JACoP plugin in Fiji.

**Protein-protein interaction studies:** GST-tagged and hexahistidine-tagged proteins were expressed in *Escherichia coli* (BL21(DE3) strain) and purified by affinity chromatography. GST-tagged proteins were purified using glutathione-Sepharose beads and hexahistidine-tagged IP6K1 and RAD51 were purified using TALON metal affinity resin (Takara Bio,

635501). For direct binding experiments, ~0.1 μM of each GST-tagged protein immobilised on glutathione-Sepharose beads was incubated with ~0.1 μM purified hexahistidine-tagged IP6K1 or RAD51 in ice-cold binding buffer (50 mM HEPES-KOH pH 7.4, 100 mM NaCl, 5% glycerol, 2 mM DTT, 5 mM $MgCl_2$, and 0.5% Triton X-100 containing protease inhibitor cocktail) at 4°C for 1 h with end-over-end mixing. Unbound proteins were removed by washing the glutathione-Sepharose beads with washing buffer (50 mM HEPES-KOH pH 7.4, 100 mM NaCl, 1% Triton X-100). To study the binding of proteins expressed in mammalian cells with RAD51 or BRCA2-CTD, HEK293T or U-2 OS cells were lysed in lysis buffer A (50 mM HEPES-KOH pH 7.4, 150 mM NaCl, 1 mM EDTA, 0.5% Nonidet P-40 with protease and phosphatase inhibitor cocktails). Clarified lysates were allowed to interact with 2-3 μg of either GST-RAD51 or GST-BRCA2-CTD immobilised on glutathione-Sepharose beads for 2 h at 4°C. Unbound proteins were removed by washing the beads thrice in wash buffer (50 mM HEPES-KOH pH 7.4, 150 mM NaCl, 1mM EDTA, 1% Nonidet P-40). To monitor the interaction of RAD51 with BRCA2-CTD in the presence of inositol phosphate analogs, 3 μg of GST-tagged RAD51 bound to glutathione-Sepharose beads was either left unphosphorylated or pre-phosphorylated with CK2 and PLK1 in the presence of $Mg^{2+}$-ATP (400 μM) in protein kinase buffer (50 mM Tris-HCl, pH 7.5, 15 mM $MgCl_2$, 0.1 mM EDTA, 1 mM EGTA, 2 mM DTT). Subsequently, beads were incubated with 50 μM 5-$InsP_7$, 5-PCP-$InsP_5$, or $InsP_6$ at 37°C for 45 min, washed in lysis buffer B (50 mM HEPES-KOH pH 7.4, 100 mM NaCl, 0.5% Nonidet P-40), and incubated overnight at 4°C with lysates from HEK293T cells overexpressing BRCA2-CTD-c-Myc (prepared in lysis buffer B) at 4°C.

For immunoprecipitation and pulldown experiments, HEK293T cells were lysed for 1 h at 4˚C in lysis buffer A. For immunoprecipitation, the lysates were incubated with the specific antibody overnight at 4˚C with end-over-end rotation, and the complexes were pulled down using either Protein A or G Sepharose beads (pre-equilibrated in lysis buffer A) for 1-2 h at 4˚C. In the case of BRCA2-CTD-c-Myc immunoprecipitation, cells were lysed in 50 mM HEPES-KOH pH 7.4, 100 mM NaCl, 10 mM EDTA, 0.5% Nonidet P-40 with protease and phosphatase inhibitor cocktails. In the case of SFB-tagged proteins, Streptavidin-Sepharose beads pre-equilibrated in lysis buffer A were added to the cell lysate, and incubated for 2 h at 4˚C. The beads were washed thrice with lysis buffer, and bound proteins were eluted by boiling the beads in 2X Laemmli buffer. The samples were processed using standard western blotting techniques, and chemiluminescence was detected using the UVITEC Alliance Q9 documentation system, or the GE ImageQuant LAS 500 imager. Densitometry analysis of

Western blots was done using Fiji software [3]. GST-tagged proteins were visualised by Ponceau S staining of the PVDF membrane.

**Mass spectrometry identification of phosphosites:** To identify phosphorylated residues on IP6K1 under three different conditions, 4 x 100 mm dishes of HEK293T cells were transfected with plasmid encoding SFB-IP6K1 (4 μg/dish) for each treatment set. Two sets of cells were treated with 0.5 μM mitomycin C (MMC) for 18 h, and one of these were allowed to recover for 12 h after removal of MMC. One set of cells was untreated. Cells were collected by scraping in chilled PBS, and lysed in lysis buffer C (20 mM Tris-HCl pH 8.0, 100 mM NaCl, 1 mM EDTA, 0.5% Nonidet P-40, 5 mM sodium fluoride, 2 mM sodium orthovanadate, 1 mM PMSF, with protease and phosphatase inhibitor cocktails) for 1 h at 4°C. Clarified lysates were incubated with 75 μL Streptavidin-Sepharose beads (pre-equilibrated in lysis buffer C) overnight at 4°C with end-over-end mixing, followed by washing and boiling the beads in 2X Laemmli buffer. Samples were resolved on a 12% SDS polyacrylamide gel, and stained with 0.1% Coomassie Brilliant Blue R-250 for 20 min. The gel was destained and the specific bands were excised and were sent to Taplin Biological Mass Spectrometry Facility (TMSF), Harvard Medical School, Boston, USA, for phosphosite identification as described earlier [4]. Gel pieces were subjected to a modified in-gel trypsin digestion procedure [5] with sequencing-grade trypsin (Promega). The extracted peptides were resolved by nano-scale reverse-phase HPLC on a microcapillary (length ~30 cm, 100 μm inner diameter) packed with 2.6 μm C18 spherical silica beads, using a gradient of 2.5-97.5% acetonitrile in 0.1% formic acid. Each eluted peptide was subjected to electron spray ionization (ESI) and allowed to enter an LTQ Orbitrap Velos Pro ion-trap mass spectrometer (Thermo Fisher Scientific). The eluted peptides were detected, isolated, and fragmented to produce a tandem mass spectrum for each peptide. The acquired fragmentation pattern was matched to translated nucleotide or protein databases using SEQUEST (Thermo Finnigan) to determine the corresponding peptide sequence. The modification of 79.9663 mass units to Ser, Thr, and Tyr was included in the database searches to identify phosphopeptides. Phosphorylation assignments (Supplementary Table S7) were determined by the Ascore algorithm [6]. According to instructions from TMSF, a phosphopeptide with an Ascore >19 was considered to be phosphorylated with 99% certainty. Mass spectrometry data were submitted to the MassIVE repository, a full member of the Proteome Xchange Consortium. Data can be accessed via the URL <https://massive.ucsd.edu>, with the data set identifier MassIVE MSV000096311

**Supplementary Table Legends**

Table S1- DNA damage causing agents used in the study

Table S2- Protein kinases used for phosphorylation assays

Table S3- Primary and secondary antibodies used in the study

Table S4- Plasmids used in the study

Table S5- Primers used in the study

Table S6- shRNA and sgRNA sequences used in the study

Table S7- Mass spectrometry-based phosphosite mapping of IP6K1

**Supplementary Figures with Legends**

Supplementary Figure S1

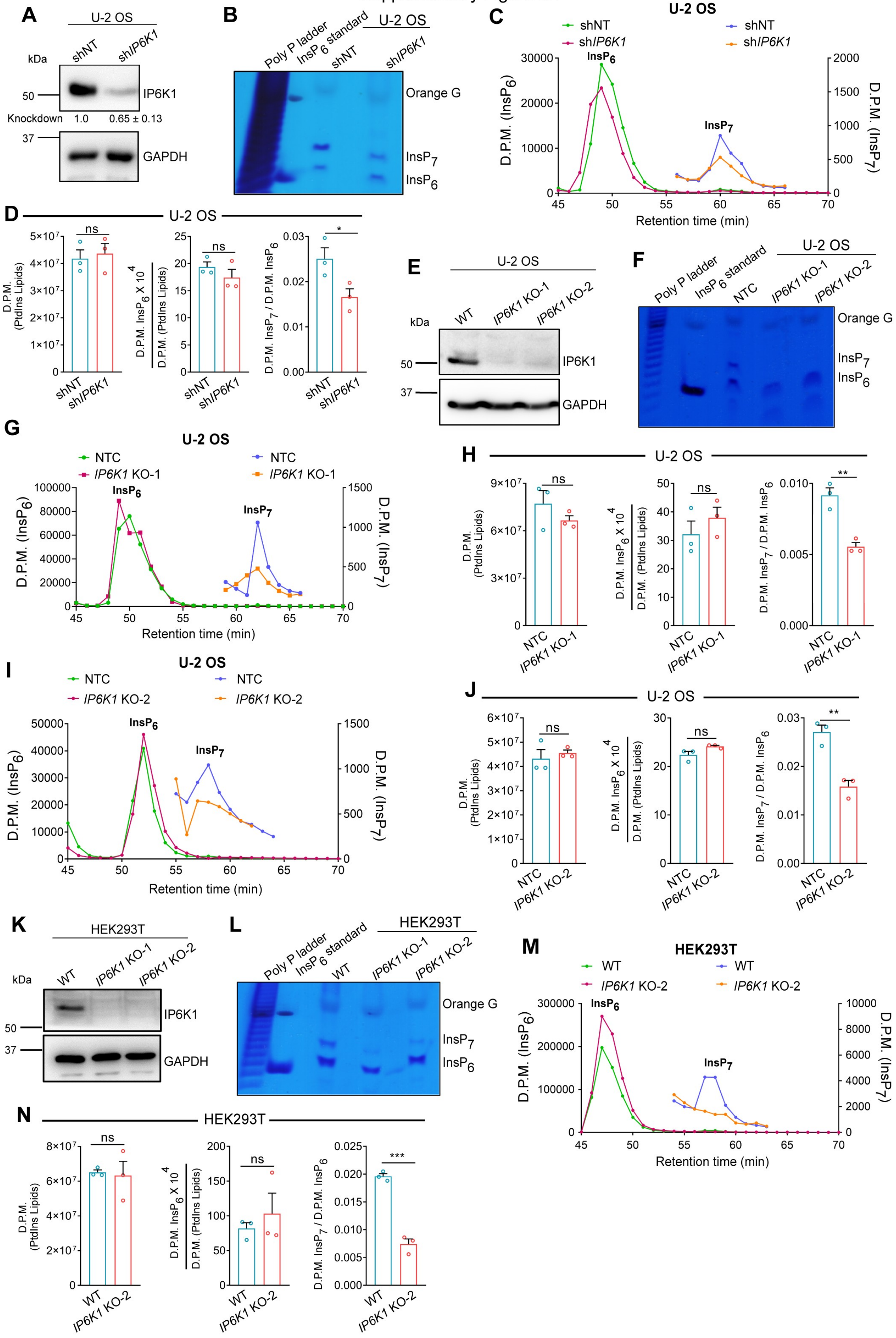

**Figure S1. Characterization of IP6K1 depleted mammalian cell lines. (A)** Representative immunoblots to detect endogenous IP6K1 in U-2 OS cells stably expressing either non-targeting control shRNA (shNT), or shRNA directed against human IP6K1 (sh*IP6K1*). GAPDH was used as a loading control. Data (mean ± SEM, $N = 4$) show the extent of IP6K1 knockdown quantified by normalising the band intensities of IP6K1 to GAPDH. **(B)** Inositol polyphosphates ($InsP_6$ and $InsP_7$) from U-2 OS shNT and sh*IP6K1* cells enriched using titanium dioxide ($TiO_2$) beads, resolved by PAGE and stained with toluidine blue. Synthetic polyphosphate (polyP) was used as a marker, purified $InsP_6$ was used as a standard, and Orange G was used in the loading dye. **(C)** Chromatogram traces depicting the SAX-HPLC analysis of [$^3$H]-inositol-labelled U-2 OS shNT and sh*IP6K1* cells showing peaks for $InsP_6$ and $InsP_7$. The traces for $InsP_7$ are also shown separately as an inset (right Y axis). **(D)** Data (mean ± SEM, $N = 3$) show the levels of phosphatidylinositol (PtdIns) lipids, $InsP_6$ normalised to PtdIns lipids, and the ratio of $InsP_7$ to $InsP_6$, for U-2 OS shNT and sh*IP6K1* cells, analyzed using a two-tailed unpaired Student's *t* test; ns, non-significant ($P > 0.05$), *$P \leq 0.05$. **(E)** Representative immunoblots to detect endogenous IP6K1 in non-targeted control (NTC) U-2 OS cells and two clonal lines of *IP6K1* knockout (KO) U-2 OS cells. GAPDH was used as a loading control. **(F)** $TiO_2$-based enrichment and PAGE analysis of $InsP_6$ and $InsP_7$ from U-2 OS NTC and *IP6K1* KO cells, as in Supplementary Figure S1B; ns, non-significant ($P > 0.05$), **$P \leq 0.01$. **(G-J)** Chromatogram traces depicting the SAX-HPLC analysis of [$^3$H]-inositol-labelled U-2 OS NTC and *IP6K1* KO-1 cells (G), and U-2 OS NTC and *IP6K1* KO-2 cells (I). Data (mean ± SEM, $N = 3$) analyzed, as in Supplementary Figure S1D, for U-2 OS NTC and *IP6K1* KO-1 (H), and U-2 OS NTC and *IP6K1* KO-2 (J) cell lines. **(K)** Representative immunoblots to detect endogenous IP6K1 in WT HEK293T cells and two clonal lines of *IP6K1* knockout (KO) HEK293T cells. GAPDH was used as a loading control. **(L)** $TiO_2$-based enrichment and PAGE analysis of $InsP_6$ and $InsP_7$ from HEK293T WT and *IP6K1* KO cell lines, as in Supplementary Figure S1B. **(M)** Chromatogram traces depicting the SAX-HPLC analysis of [$^3$H]-inositol-labelled HEK293T WT and *IP6K1* KO-2 cell lines. The comparison of HEK293T WT and *IP6K1* KO-1 cell lines has been presented in [2]. **(N)** Data (mean ± SEM, $N = 3$) analyzed, as in Supplementary Figure S1D, for HEK293T WT and *IP6K1* KO-2 cell lines; ; ns, non-significant ($P > 0.05$), ***$P \leq 0.001$.

Supplementary Figure S2

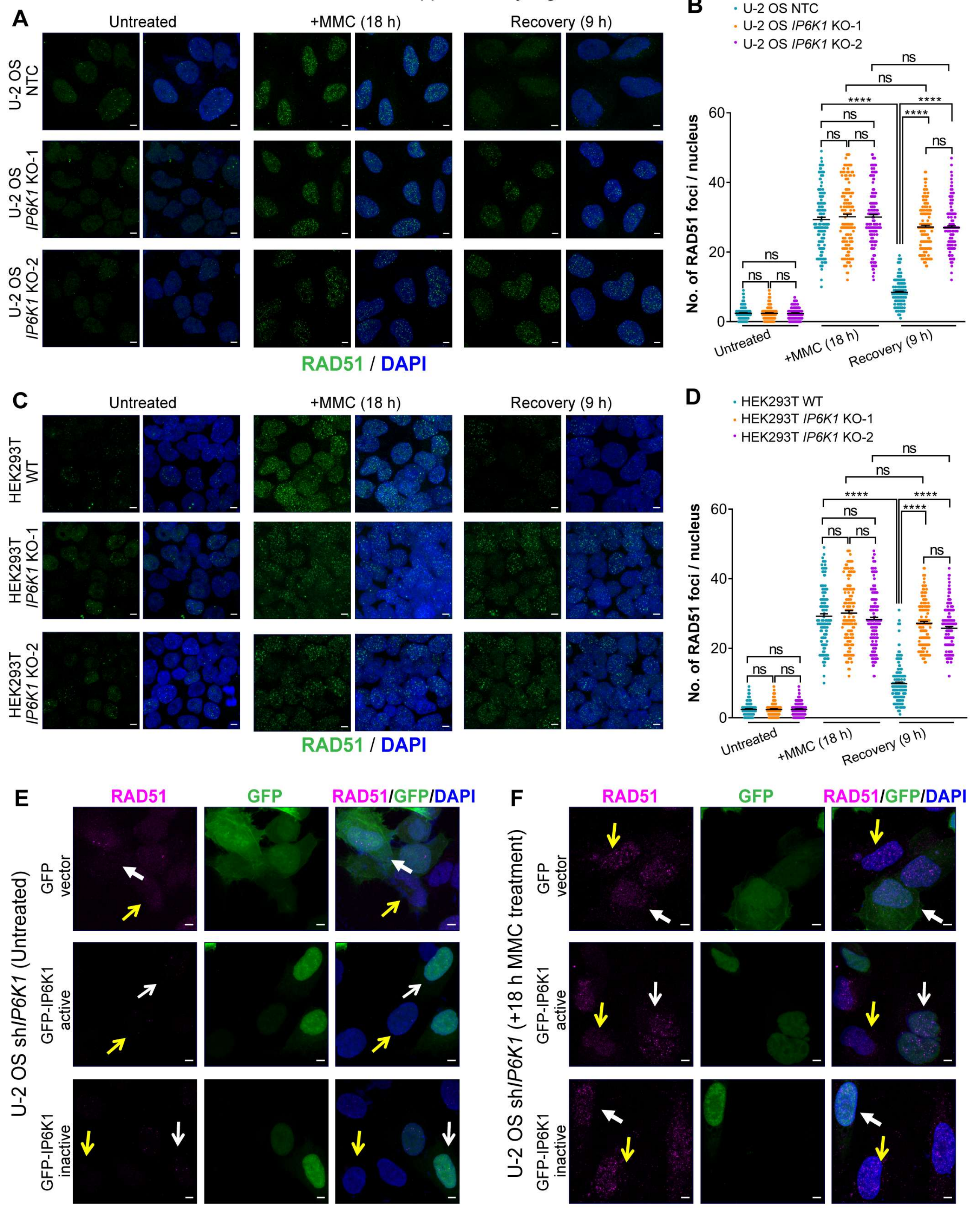


**Figure S2. MMC-induced RAD51 foci persist in *IP6K1* KO cells. (A)** Representative immunofluorescence images of NTC U-2 OS cells and two clonal lines of *IP6K1* knockout (KO) U-2 OS cells, left untreated, treated with MMC, or allowed to recover from MMC-induced DNA damage, and stained with anti-RAD51 antibody (green). **(B)** Quantification of the number of RAD51 foci per nucleus in (A). Data distribution is represented by scatter dot plots ($n$ = 151, 160, and 169 nuclei for untreated, MMC treated and post-recovery NTC U-2 OS cells; $n$ = 142, 145, and 151 nuclei for untreated, MMC treated and post-recovery *IP6K1* KO-1 U-2 OS cells; and $n$ = 152, 161, and 158 nuclei for untreated, MMC treated and post-recovery *IP6K1* KO-2 U-2 OS cells, from four independent experiments). **(C)** Representative immunofluorescence images of WT HEK293T and two clonal lines of *IP6K1* knockout (KO) HEK293T cells, left untreated, treated with MMC, or allowed to recover from MMC-induced DNA damage, and stained with anti-RAD51 antibody (green). **(D)** Quantification of the number of RAD51 foci per nucleus in (C). Data distribution is represented by scatter dot plots ($n$ = 151, 158, and 169 nuclei for untreated, MMC treated and post-recovery WT HEK293T cells; $n$ = 142, 152, and 151 nuclei for untreated, MMC treated and post-recovery *IP6K1* KO-1 HEK293T cells; and $n$ = 147, 158, and 150 for untreated, MMC treated and post-recovery *IP6K1* KO-2 HEK293T cells, from four independent experiments). Data presented in B, and D were analyzed using Kruskal-Wallis test with Dunn's multiple comparison post hoc test; ns, non-significant ($P > 0.05$), ****$P \leq 0.0001$. **(E and F)** Representative immunofluorescence images of U-2 OS sh*IP6K1* cells overexpressing GFP (green), GFP-tagged active mouse IP6K1, or inactive (K226A/S334A) mouse IP6K1, left untreated (E) or treated with MMC (F), and stained with anti-RAD51 antibody (magenta). Transfected and untransfected cells are indicated with white and yellow arrows respectively. Nuclei were stained with DAPI (blue), and scale bars are 5 μm in A, C, E, and F.

Supplementary Figure S3

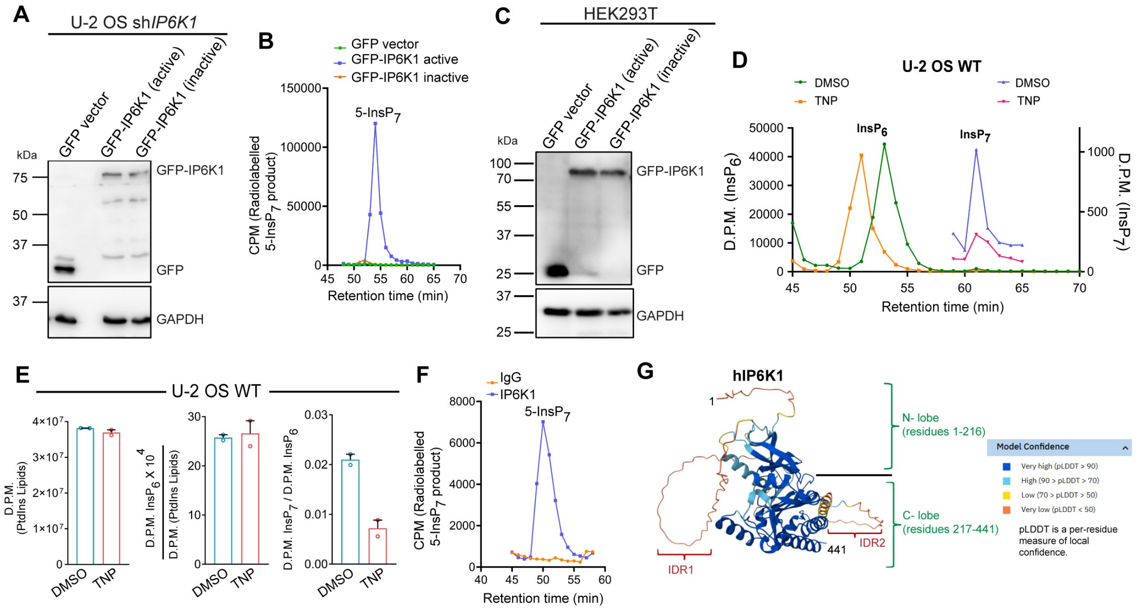


**Figure S3. Analysis of IP6K1 enzyme activity. (A)** Representative immunoblots to detect GFP (control), GFP-tagged active mouse IP6K1, and GFP-tagged inactive (K226A/S334A) mouse IP6K1 overexpressed in U-2 OS sh*IP6K1* cells. GAPDH was used as a loading control. **(B)** Confirmation of loss of IP6K1 activity in K226A/S334A mutant. Chromatogram traces depicting the SAX-HPLC analysis of 5[β-$^{32}$P]InsP$_7$ formed during the incubation of immunoprecipitated GFP, active GFP-IP6K1, or inactive GFP-IP6K1 with [γ-$^{32}$P]ATP and InsP$_6$. **(C)** The expression levels of GFP, active GFP-IP6K1, and inactive GFP-IP6K1 overexpressed in HEK293T cells. Lysates from these cells were used to assess IP6K1 activity shown in (B). **(D)** Chromatogram traces depicting the SAX-HPLC analysis of [$^3$H]-inositol-labelled WT U-2 OS cells treated with either DMSO (vehicle control) or TNP (10 μM) for 60 h, showing peaks for InsP$_6$ and InsP$_7$. The traces for InsP$_7$ are also shown separately as an inset (right Y axis). **(E)** Data (mean ± range, $N = 2$) show the levels of PtdIns lipids, InsP$_6$ normalised to PtdIns lipids, and the ratio of InsP$_7$ to InsP$_6$, WT U-2 OS cells treated with either DMSO or TNP. **(F)** Chromatogram traces depicting the SAX-HPLC analysis of 5[β-$^{32}$P]InsP$_7$ formed during the incubation of endogenous IP6K1 immunoprecipitated from HEK293T cells with [γ-$^{32}$P]ATP and InsP$_6$. Normal rabbit IgG incubated with HEK293T cell lysate served as a control. **(G)** The structure of human IP6K1, as predicted by AlphaFold Monomer v2.0 [7]. The color key for the AlphaFold pLDDT score, used as the per residue confidence metric, is indicated. Intrinsically disordered regions (IDRs), marked IDR1 and IDR2 within the N-lobe and C-lobes respectively, are shown as low/very low confidence regions in the predicted structure.

Supplementary Figure S4

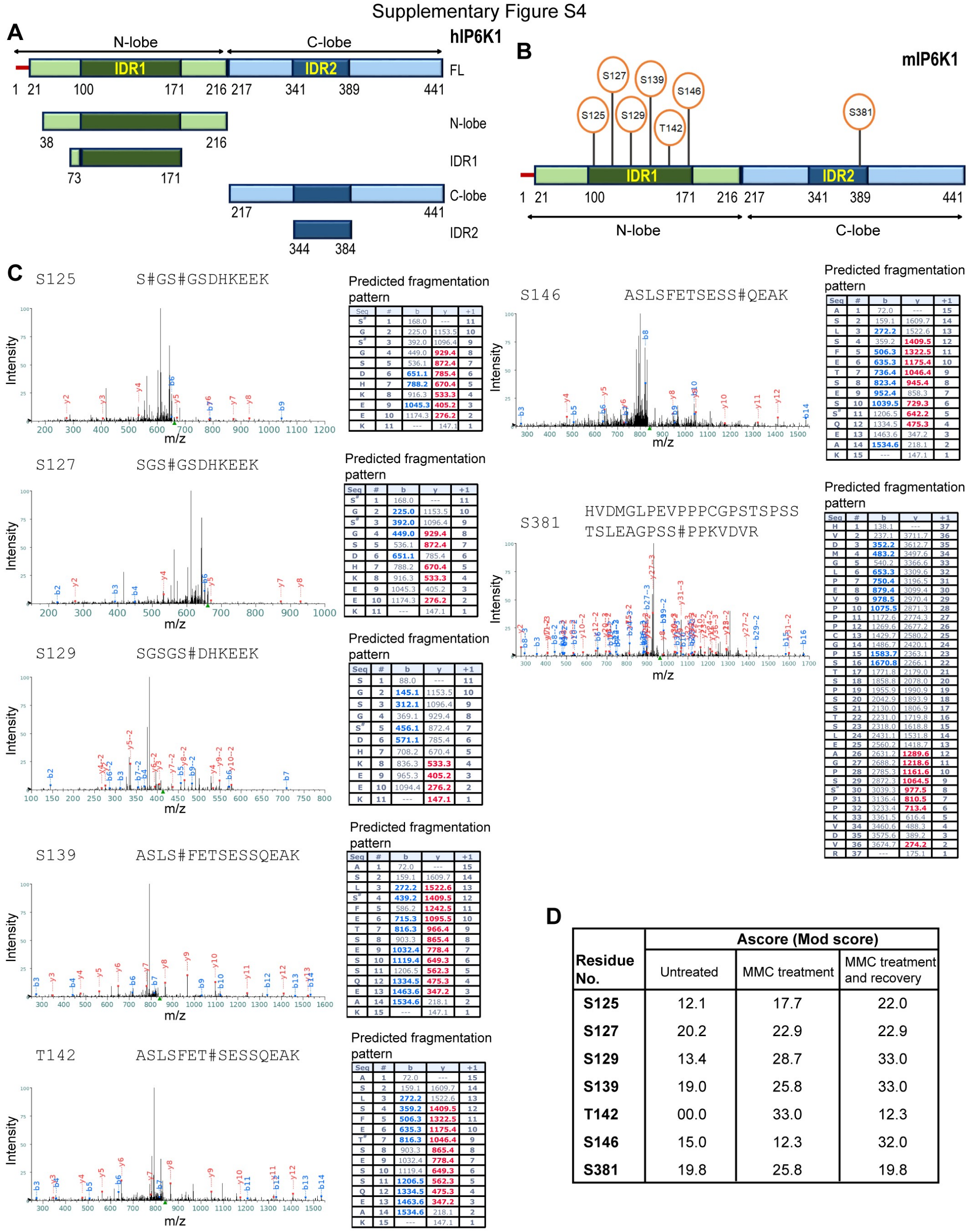


| Residue No. | Ascore (Mod score) Untreated | MMC treatment | MMC treatment and recovery |
|---|---|---|---|
| S125 | 12.1 | 17.7 | 22.0 |
| S127 | 20.2 | 22.9 | 22.9 |
| S129 | 13.4 | 28.7 | 33.0 |
| S139 | 19.0 | 25.8 | 33.0 |
| T142 | 00.0 | 33.0 | 12.3 |
| S146 | 15.0 | 12.3 | 32.0 |
| S381 | 19.8 | 25.8 | 19.8 |

**Figure S4. Phosphosite analysis on IP6K1 by mass spectrometry. (A)** Schematic representation of predicted domain map of human IP6K1 based on disorder prediction using IUPred3 (https://iupred3.elte.hu/) and tertiary structure prediction using AlphaFold Monomer v2.0 (https://alphafold.ebi.ac.uk/entry/AF-Q92551-F1). Human IP6K1 is divided into an N-lobe and a C-lobe, each encompassing intrinsically disordered regions, IDR1 and IDR2, respectively. The amino acid residue numbers corresponding to each lobe and IDRs are marked. The GST-fusion constructs corresponding to IP6K1 fragments used in Figure 4A and B are indicated. **(B)** Schematic (as in A) indicating the specific residues on IP6K1 detected to be phosphorylated by mass spectrometry analysis of SFB-tagged mouse IP6K1 overexpressed in HEK293T cells under untreated, MMC treated, and post-recovery conditions. **(C)** Representative MS/MS spectra identifying the indicated phosphorylated residues on overexpressed SFB-IP6K1. The sequence of the tryptic fragment is shown above each spectrum, and the hash symbol (#) to the right of the amino acid indicates the modified residue. The masses of the b ions (blue) and y ions (red) detected in the peptide fragmentation pattern are shown in the table to the right. **(D)** The table provides the statistical confidence of phosphosite assignment (Ascore) at the indicated residues on IP6K1 under each treatment condition, as detailed in Supplementary Table S7. A site with an Ascore >19 was considered to be phosphorylated with 99% certainty.

Supplementary Figure S5

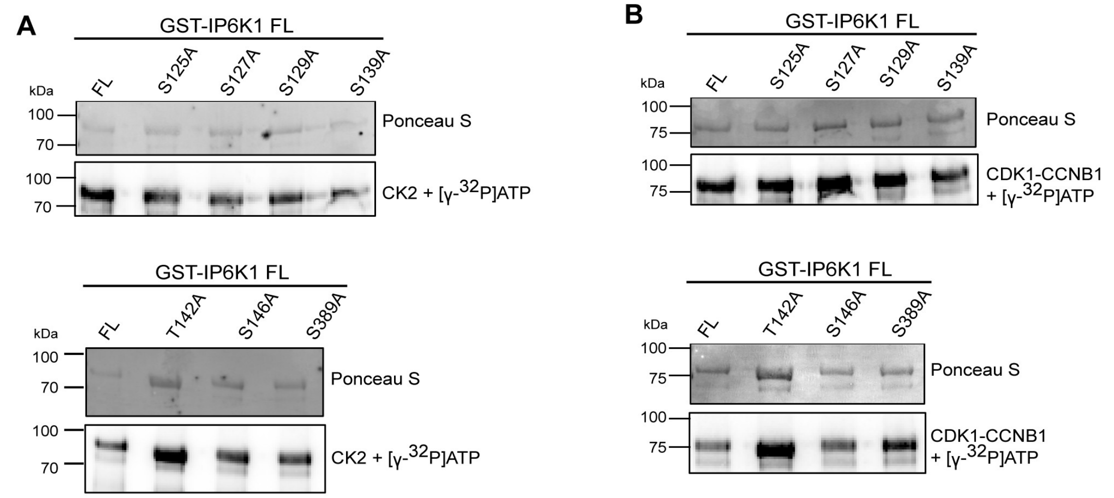


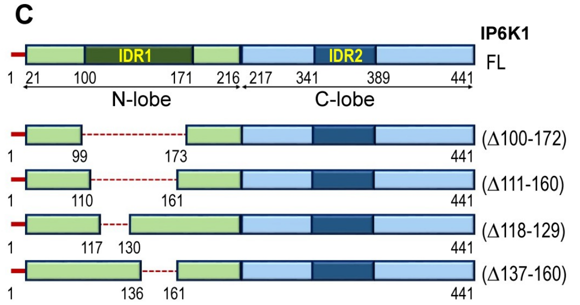


**D**

**>sp|Q92551|IP6K1_HUMAN**

**IP6K1(Δ100-172)**

**Δ[100** ETVEQDDTTEREQPRRKHSRRSLHRSGS
GSDHKEEKASLSLETSESSQEAKSPKVE
LHSHSEVPFQMLDGNSG **172]**

**IP6K1(Δ111-160)**

**Δ[111** EQPRRKHSRRSLHRSGSGSDHKEEKAS
LSLETSESSQEAKSPKVELHSHS **160]**

**IP6K1(Δ118-129)**

**Δ[118** SRRSLHRSGSGS **129]**

**IP6K1(Δ137-160)**

**Δ[137** SLSLETSESSQEAKSPKVELHSHS **160]**

**Figure S5. IDR1 mutants in IP6K1. (A and B)** Ser/Thr to Ala point mutant forms of full length (FL) GST-tagged human IP6K1 (GST-IP6K1) expressed in *E. coli* were subjected to phosphorylation by either CK2 (A) or CDK1-CCNB1 (B) in presence of [$\gamma$-$^{32}$P]ATP. Representative images show autoradiography to detect phosphorylation mediated by the protein kinases (right) and GST-tagged proteins stained with Ponceau S (left) ($N$ = 2). **(C)** Schematic representation of in-frame IDR1 deletion mutants of human IP6K1 used in Figure 4C - F. The dotted line in each mutant depicts the deleted region, and the amino acid residues that mark the boundary of the deleted fragments are indicated on the bar diagrams. **(D)** The amino acid sequences of deleted regions in human IP6K1 shown in (C).

Supplementary Figure S6

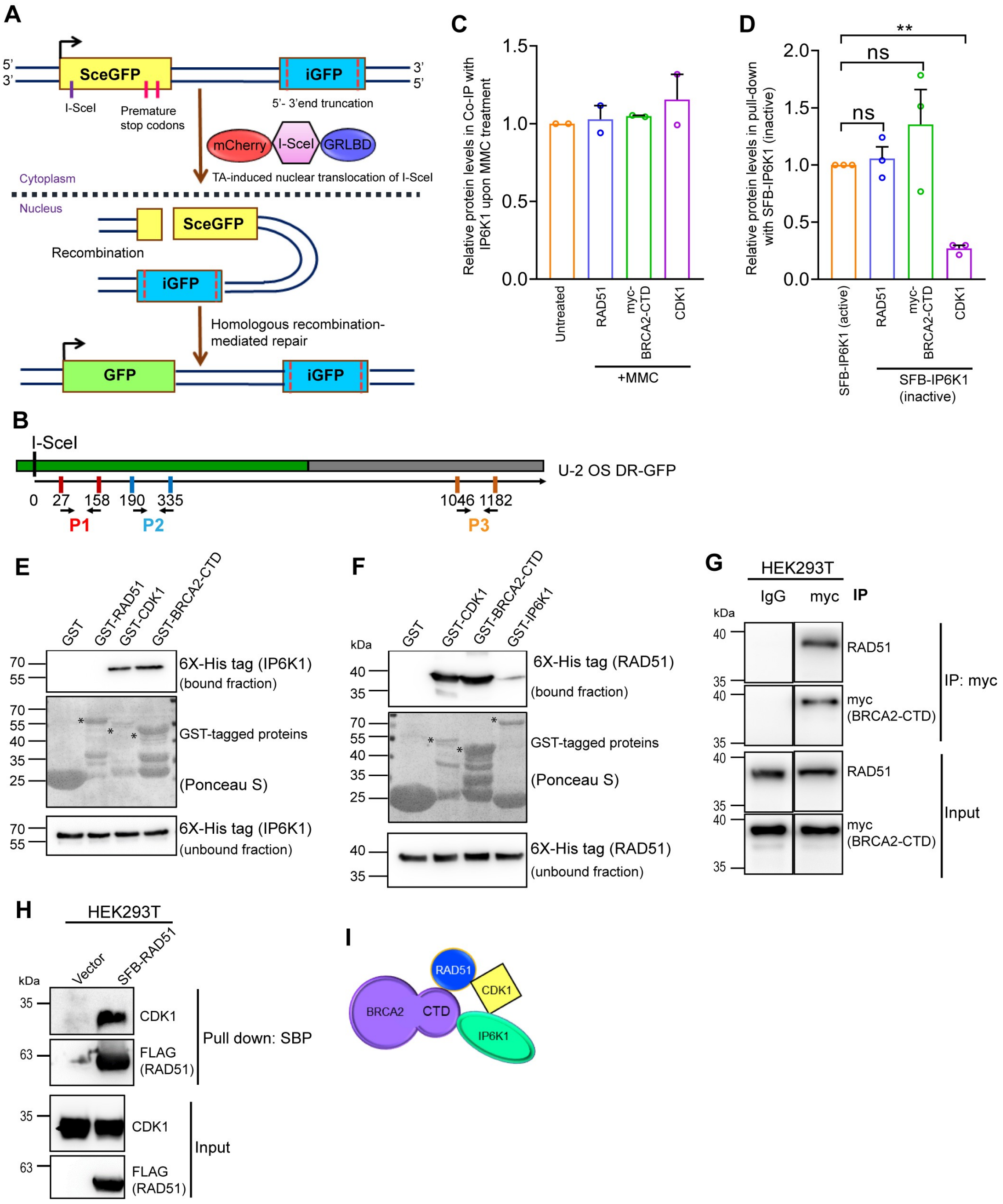


**Figure S6. HR reporter assay and protein-protein interactions. (A)** Schematic representation of the U-2 OS DR-GFP (mCherry-I-SceI-GR) HR reporter system. The reporter is composed of two mutant copies of GFP, one with a promoter and two premature stop codons that also harbors a cleavage site for the I-Sce-I endonuclease (SceGFP), and a second promoterless sequence that has upstream and downstream truncations rendering it inactive (iGFP). cDNA encoding I-SceI fused with the glucocorticoid receptor ligand binding domain

(GRLBD) and mCherry is stably integrated in this U-2 OS reporter line. Addition of the glucocorticoid receptor agonist triamcinolone acetonide (TA), which binds to GRLBD, leads to nuclear translocation of I-SceI, and endonucleolytic cleavage at SceGFP. The double strand break is repaired by HR, using iGFP as the template, generating a sequence encoding active GFP, which is assessed by flow cytometry. **(B)** Schematic indicating the locations of three sets of primers (P1, P2, and P3) used in the ChIP-qPCR assay shown in Figure 5C. The numbers mark the distance (in base pairs) from the I-SceI cleavage site corresponding to the ends of the fragment amplified during qPCR. The genomic region where HR is expected is marked in green. **(C)** Quantification of Figure 5D - F, showing the interaction of endogenous IP6K1 in HEK293T cells with endogenous RAD51, overexpressed myc-tagged BRCA2-CTD, and endogenous CDK1. Data (mean ± range, $N = 2$ for RAD51; $N = 2$ myc-BRCA2-CTD; and mean ± SEM, $N = 2$ for CDK1) show the extent of interaction of each protein with IP6K1 in MMC treated cell lysates, normalised to the respective interaction in untreated cells. **(D)** Quantification of Figure 5G - I, showing the interaction of active or inactive (K226A/S334A) SFB-tagged mouse IP6K1 overexpressed in HEK293T cells with endogenous RAD51, myc-BRCA2-CTD, and endogenous CDK1. Data (mean ± SEM, $N = 3$ for RAD51; $N = 3$ for myc-BRCA2-CTD; and $N = 3$ for CDK1) show the extent of interaction of each protein with inactive (K226A/S334A) IP6K1 normalised to the respective interaction with active IP6K1, and were analyzed using a one-sample *t* test; ns, non-significant ($P > 0.05$), $^{**}P \leq 0.01$. **(E and F)** Representative immunoblots examining binding of the indicated GST-tagged proteins with hexahistidine-tagged mouse IP6K1 (E), or hexahistidine-tagged human RAD51 (F). The interactions were detected by immunoblotting with anti-6X His tag antibody, and GST-tagged proteins were visualised by staining with Ponceau S (N = 3). The asterisks (*) mark specific bands corresponding to the GST-tagged proteins. **(G)** Representative immunoblots examining co-immunoprecipitation of endogenous RAD51 with overexpressed myc-BRCA2-CTD in HEK293T lysates from MMC treated cells. IgG was used as a control. **(H)** Representative immunoblots examining pull-down (PD) of endogenous CDK1 with overexpressed SFB-RAD51 in lysates from MMC treated HEK293T cells. SFB-RAD51 was pulled down with Streptavidin Sepharose beads (which bind the Streptavidin-binding peptide, SBP) and probed with an anti-FLAG antibody. **(I)** Illustration depicting the direct or indirect interaction of the indicated proteins with IP6K1 upon MMC treatment.

Supplementary Figure S7

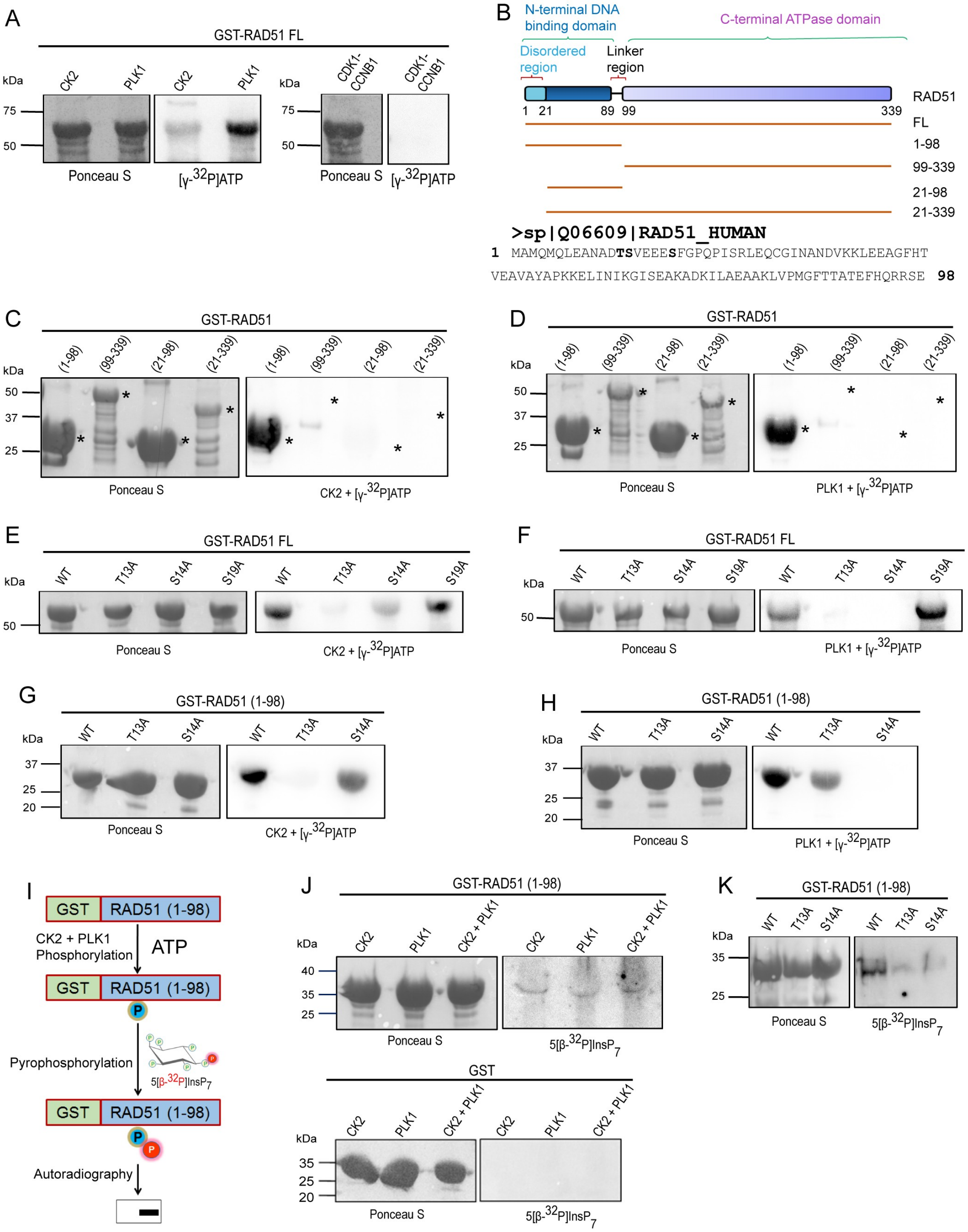


**Figure S7. 5-InsP$_7$ mediated pyrophosphorylation of RAD51. (A)** GST-tagged full-length RAD51 (GST-RAD51 FL) expressed in *E. coli* was incubated with CK2, PLK1, or CDK1-CCNB1 in presence of [γ-$^{32}$P]ATP. Representative images show autoradiography to detect phosphorylation mediated by protein kinases (right) and GST-RAD51 was detected by Ponceau S staining (left) ($N$ = 2). **(B)** Schematic representation of domain map of human

RAD51 and fragments tested for phosphorylation in this study. The protein sequence of the N-terminal DNA binding domain of RAD51 is shown, with the tested phosphorylation sites shown in bold. **(C and D)** The GST-tagged RAD51 fragments shown in (B) expressed in *E. coli* were incubated with either CK2 (C) or PLK1 (D) in presence of radiolabelled ATP ([γ-$^{32}$P]ATP) ($N$ = 2). **(E -H)** The indicated Ser/Thr to Ala mutant forms of GST-RAD51 FL and fragment (1-98) expressed in *E. coli* were incubated with CK2 (E and G) or PLK1 (F and H) in presence of radiolabelled ATP ([γ-$^{32}$P]ATP) ($N$ = 2). **(I)** Schematic depicting the method to test 5-$InsP_7$-mediated pyrophosphorylation of RAD51. GST-RAD51 (1-98) expressed in *E. coli* and immobilized on glutathione Sepharose beads was first incubated with protein kinases CK2 and/or PLK1 in the presence of unlabelled ATP, the beads were washed, and subsequently incubated with 5[β-$^{32}$P]$InsP_7$, followed by autoradiography to detect pyrophosphorylation. **(J)** Pre-phosphorylation GST-RAD51 (1-98) by CK2, PLK1, or both protein kinases, and subsequent pyrophosphorylation by 5[β-$^{32}$P]$InsP_7$, as described in (I). GST was used as a control. **(K)** GST-RAD51 (1-98) WT and the indicated Ser/Thr to Ala mutants were subjected to pre-phosphorylation by CK2 and PLK1 in presence of unlabelled ATP, followed by incubation with 5[β-$^{32}$P]$InsP_7$ to test their pyrophosphorylation. Images in (C-H, J, and K) show autoradiography to detect either phosphorylation or pyrophosphorylation (right) and GST-tagged proteins stained with Ponceau S (left).

Supplementary Figure S8

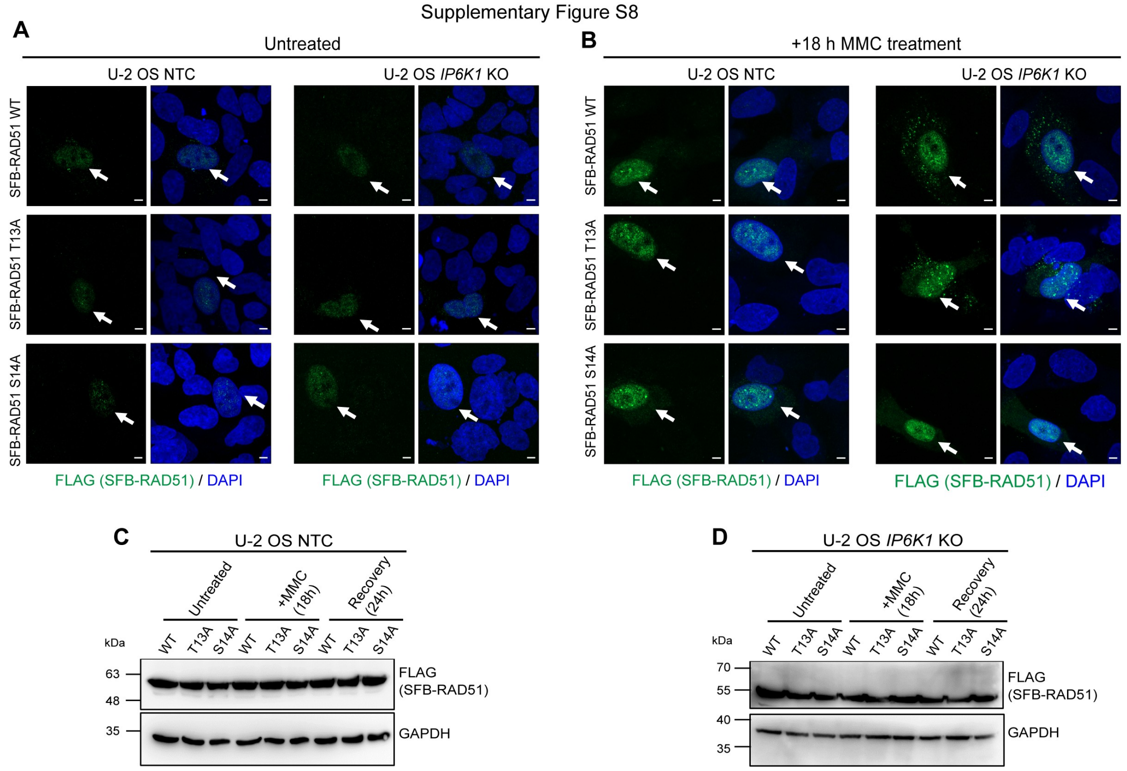


**Figure S8. Effect of MMC treatment on RAD51 mutants. (A and B)** Representative immunofluorescence images of U-2 OS NTC and *IP6K1* KO cells overexpressing SFB-tagged RAD51 WT, T13A, or S14A, that were left untreated (A), or treated with MMC (B), stained with anti-FLAG antibody (green) to detect the SFB-tag. Transfected cells are indicated with white arrows, nuclei were stained with DAPI (blue), and scale bars are 5 µm. **(C and D).** Representative immunoblots of lysates from U-2 OS NTC (C) and *IP6K1* KO (D) cell lines overexpressing SFB-RAD51 WT, T13A, or S14A, that were left untreated, treated with MMC, or allowed to recover from MMC-induced DNA damage for the indicated time. Blots were probed with an anti-FLAG antibody to detect the SFB-tag and GAPDH served as the loading control.
